# Ringer Loss in *Drosophila* Uncovers Mitochondrial Complex I Deficits Relevant to Human Parkinson’s Disease

**DOI:** 10.64898/2026.07.27.741040

**Authors:** Haven G. Tillmon, Savannah K. Boyen, Raquel Velazquez, Michelle E. Urbina-Berlanga, Marco Sciortino, Swati Banerjee

## Abstract

Tubulin polymerization promoting proteins (TPPPs) are known for their cytoskeletal regulation across species; however, emerging evidence suggests broader cellular functions, including potential roles in mitochondrial biology. Here, we identify the *Drosophila* homolog of human TPPP, Ringer, as a previously unrecognized regulator of mitochondrial bioenergetics and electron transport chain Complex I (CI) function. Ringer is enriched in the mitochondrial matrix, and its loss results in reduced levels of multiple CI subunits and assembly factors and a significant decrease in CI enzymatic activity. Notably, in addition to TPPP dysfunction, similar deficits are observed in postmortem human Parkinson’s disease (PD) brain tissues, underscoring the translational relevance of our *Drosophila* model. Pharmacological administration of resveratrol, a reactive oxygen species (ROS) scavenger, reduces mitochondrial superoxide levels and improves CI enzymatic activity and ATP production in *ringer* mutants, demonstrating that targeted antioxidant therapeutics can improve bioenergetic functions in *ringer* mutants. Together, these findings establish Ringer as a key regulator of mitochondrial bioenergetics and reveal Ringer-induced CI dysfunction as a phenotype relevant to mitochondrial abnormalities observed in PD, and provide a robust and translationally meaningful framework for investigating mitochondrial dysfunction in PD.

## INTRODUCTION

Tubulin polymerization promoting proteins (TPPPs) are a family of highly conserved microtubule-associated proteins (MAPs) essential for microtubule polymerization, stabilization, and bundling.^1–4^ Although originally characterized as predominantly cytosolic, subsequent studies have demonstrated that TPPPs are found in additional subcellular compartments, including the nucleus^5^ and mitochondria,^6^ suggesting broader regulatory functions beyond cytoskeletal organization. Growing evidence suggests that TPPPs influence a wide array of cellular activities, extending far beyond their canonical roles in microtubule regulation and encompassing multiple aspects of cellular homeostasis.^7^

Three TPPPs exist in humans (TPPP1, TPPP2, and TPPP3), and the *Drosophila* protein Ringer shows high sequence homology and function to TPPP1 and TPPP3, reflecting its conserved structural and functional roles.^1,5^ TPPP1 is a 25-kDa intrinsically disordered protein with particularly high conservation in its C-terminal region, where the last 50 amino acids form the most conserved domain across species.^8,9^ In the mammalian nervous system, TPPP1 is enriched in the brain and expressed at higher levels in oligodendrocytes^10^ and is critical for myelination.^11^ In *Drosophila*, the TPPP homolog Ringer contributes to neuronal health by regulating axon outgrowth and regeneration,^12^ as well as synaptic growth and neurotransmission.^3,4,6^ Importantly, human TPPP1 is implicated in α-synucleinopathies, including Parkinson’s disease (PD), yet its involvement in PD pathogenesis and/or progression remains incompletely understood.^13–15^

Mitochondrial dysfunction, particularly impairment of electron transport chain complex I (CI), is a well-established hallmark of PD and is among the more frequently reported biochemical abnormalities in patient tissues,^16,17^ toxin-based models,^18–22^ and genetic systems.^23,24^ CI deficits contribute to elevated ROS generation, impaired oxidative phosphorylation, and heightened vulnerability to metabolic stress, all of which converge on dopaminergic neuron degeneration in PD.^16,25,26^ Mitochondrial toxins such as 1-methyl-4-phenyl-1,2,3,6-tetrahydropyridine (MPTP)^18,19^ and rotenone,^20,21^ which selectively inhibit CI, reliably produce PD-like pathology in cellular and animal models, underscoring the central role of CI impairment in experimental systems of neurodegeneration. Despite this strong association, the upstream mechanisms destabilizing CI in PD remain insufficiently defined.

We previously identified the single *Drosophila* homolog of human TPPP, Ringer,^3^ and demonstrated that loss of Ringer recapitulates many salient features of human PD, including progressive locomotor deficits, neurodegeneration, and reduced lifespan in adult *ringer* mutants.^6^ Among additional pathological correlates, *ringer* mutants displayed robust mitochondrial structural damage and dysfunction, including increased superoxide levels, decreased mitochondrial membrane potential, and reduced ATP levels.^6^ These mitochondrial vulnerabilities provide a critical framework for investigating how TPPP may contribute to impaired mitochondrial function and possibly disease pathogenesis.

Here, we demonstrate that Ringer is found not only in the cytoplasm, but also associates with mitochondria, where it is specifically enriched in the mitochondrial matrix, and its loss leads to pronounced mitochondrial bioenergetic deficits. Ringer loss results in decreased levels of multiple CI matrix subunits as well as CI assembly proteins. Iodixanol density gradient centrifugation followed by immunoblotting analyses of wild-type brain lysates revealed that Ringer co-sediments with CI subunits, and loss of Ringer alters this distribution. Consistent with these deficits, CI enzymatic activity is significantly reduced in *ringer* mutants. Pharmacological administration of the ROS scavenger resveratrol lowers the elevated mitochondrial ROS burden in *ringer* mutants while improving CI activity and increasing ATP production, indicating that oxidative stress contributes to mitochondrial dysfunction in *ringer* mutants. Importantly, CI-related mitochondrial abnormalities observed in *ringer* mutants are partially recapitulated in postmortem human PD brain tissue, where biochemical analyses of parietal cortex Brodmann area 40 (BA40) samples reveal reduced levels of specific CI proteins. Additionally, immunohistochemical analysis of TPPP in the BA40 brain region shows altered localization in the form of aggregates in PD patients compared to age- and sex-matched controls. Together, these findings establish *ringer* mutants as a powerful *in vivo* model for dissecting mammalian TPPP biology and provide insight into how mitochondrial CI dysfunction may contribute to PD-related mitochondrial pathology and neurodegeneration.

## RESULTS

### Ringer is highly enriched in the mitochondrial matrix

Our prior work revealed that Ringer localizes to the mitochondria and that *ringer* mutants display mitochondrial ultrastructural defects and bioenergetic dysfunction^6^. We therefore sought to determine the quantitative distribution of Ringer within the mitochondrial compartments. We first performed a subcellular fractionation to obtain the mitochondrial (Mito, Fig. 1A) and cytosolic (Cyto, Fig. 1A) fractions from wild-type (+/+, Fig. 1A) and *ringer* (-/-, Fig. 1A) mutant adult brain total lysates (Total, Fig. 1A) as previously reported,^6^ followed by immunoblotting analysis. Upon quantification, we found that Ringer is equally distributed between the mitochondria and cytosol in the wild-type *Drosophila* brain with respect to the total lysates (Fig. 1B). As expected, the *ringer* mutant brain fractions (Fig. 1A) show loss of Ringer. Anti-GAPDH^27^ and anti-Porin^28^ were used to demonstrate that there was no cross-contamination between the mitochondrial and cytosolic fractions (Fig. 1A).

**Figure 1:**
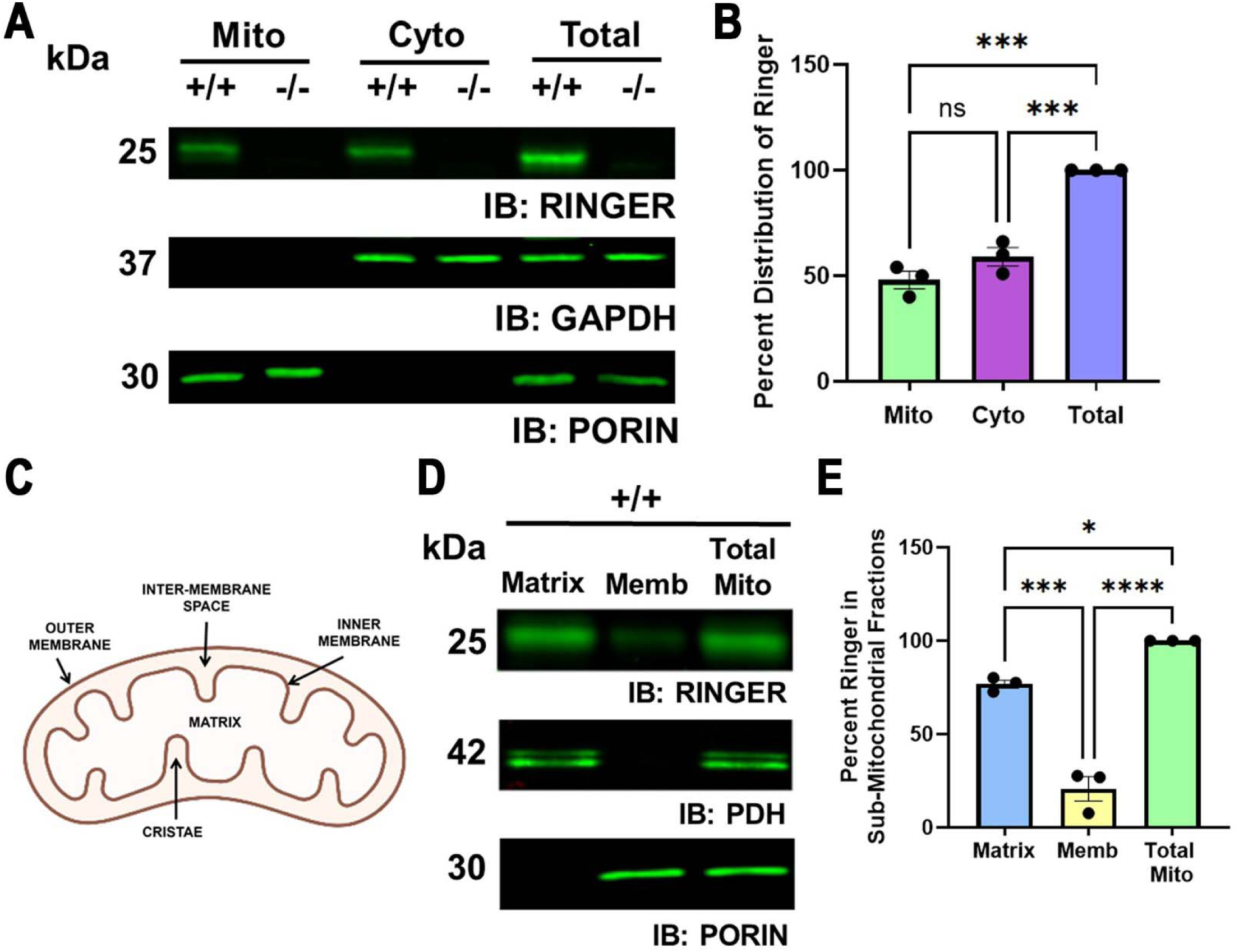
Ringer is distributed equally in both cytosolic and mitochondrial fractions and is enriched in the mitochondrial matrix. **A** Immunoblots of subcellular fractions from wild-type (+/+) and *ringer* mutant (−/−) fly heads probed with anti-Ringer (25kDa), anti-GAPDH (37kDa), and anti-Porin (30kDa) across mitochondrial (mito), cytosolic (cyto), and total lysate fractions. **B** Quantification of Ringer in mitochondrial (green) and cytosolic (magenta) fractions with respect to the total lysates (lavender) from wild-type (+/+) fly head lysates. Data are presented as mean ±SEM. Statistical significance was determined using a one-way ANOVA followed by Tukey’s multiple comparisons test. F(6)=61.16, mito vs. cyto ns p=0.1433; total vs. mito ***p=0.0001; total vs. cyto ****p=0.0004 from n=3 experiments. **C** Schematic of mitochondrial structure highlighting the outer and inner membranes, cristae, inter-membrane space, and matrix. **D** Immunoblots of sub-mitochondrial fractions from wild-type (+/+) fly heads probed with anti-Ringer (25kDa), anti-PDH (42 kDa), and anti-Porin (30kDa) across matrix, membrane (memb), and total mitochondrial fractions. **E** Quantification of Ringer band intensity in the matrix (blue) and membrane (yellow) fractions with respect to the total mitochondria (green) from wild-type (+/+) fly head lysates. Data are presented as mean ±SEM. Statistical significance was determined using a one-way ANOVA followed by Tukey’s multiple comparisons test. F(6)=103.9, total mito vs. matrx *p=0.0149; matrix vs. memb ***p=0.0001; and total mito vs. memb ****p<0.0001 from n=3 experiments.

We next determined Ringer distribution within the mitochondria (Fig. 1C) using sub-mitochondrial fractionation. Mitochondrial fractions were first isolated from the wild-type brain lysates (Total Mito, Fig. 1D) and further separated into matrix and membrane fractions using ultracentrifugation and immunoblotting (Fig. 1D). We found that Ringer is predominantly associated with the mitochondrial matrix (Fig. 1D, quantified in E) and minimally present in the mitochondrial membrane (Fig. 1D, quantified in E) with respect to the total mitochondria. The same immunoblots were probed with a marker for the mitochondrial matrix, anti-pyruvate dehydrogenase (anti-PDH),^29^ and anti-Porin was used as a marker for the outer mitochondrial membrane to assess the purity of the matrix and membrane fractions and the absence of cross-contamination between them (Fig. 1D). These results demonstrate that Ringer is associated with the mitochondria and is enriched in the mitochondrial matrix.

### Ringer loss reduces Complex I matrix and assembly proteins

Having established that Ringer is enriched in the mitochondrial matrix, we next examined whether loss of Ringer alters the protein levels of CI subunits and assembly factors (Fig 2A). We examined two of the core CI subunits from the matrix arm, NDUFS2 and NDUFS3,^30^ CI assembly proteins, NDUFAF1 and ECSIT,^31^ CI proteins from the membrane arm, NDUFB8 and NDUFB10, as well as a hinge protein, NDUFA9.^30^ We performed immunoblotting analysis of mitochondrial fractions isolated from the adult *Drosophila* brains of both wild-type (+/+) and *ringer* mutants (-/-) and probed with antibodies against NDUFS2, NDUFS3, NDUFB8, NDUFB10, NDUFAF1, NDUFA9 and ECSIT (Fig. 2B). NDUFS2 (Fig. 2C), NDUFS3 (Fig. 2D), NDUFAF1 (Fig. 2H), and ECSIT (Fig. 2I) showed significantly decreased levels in *ringer* mutants compared to wild-type controls. Immunoblotting analysis for ECSIT was performed on total lysate fractions (Fig. 2B) isolated from the *Drosophila* brain, as ECSIT was primarily detected in the cytosolic fraction (Supplemental Fig. 1A-B). Interestingly, subunits from the CI membrane arm, NDUFB8 (Fig. 2E), NDUFB10 (Fig. 2F), and hinge, NDUFA9 (Fig. 2G), did not show any significant changes in their respective levels in *ringer* mutants compared to wild-type controls. These findings suggest that loss of Ringer alters the levels of specified CI matrix arm subunits and assembly proteins.

**Figure 2:**
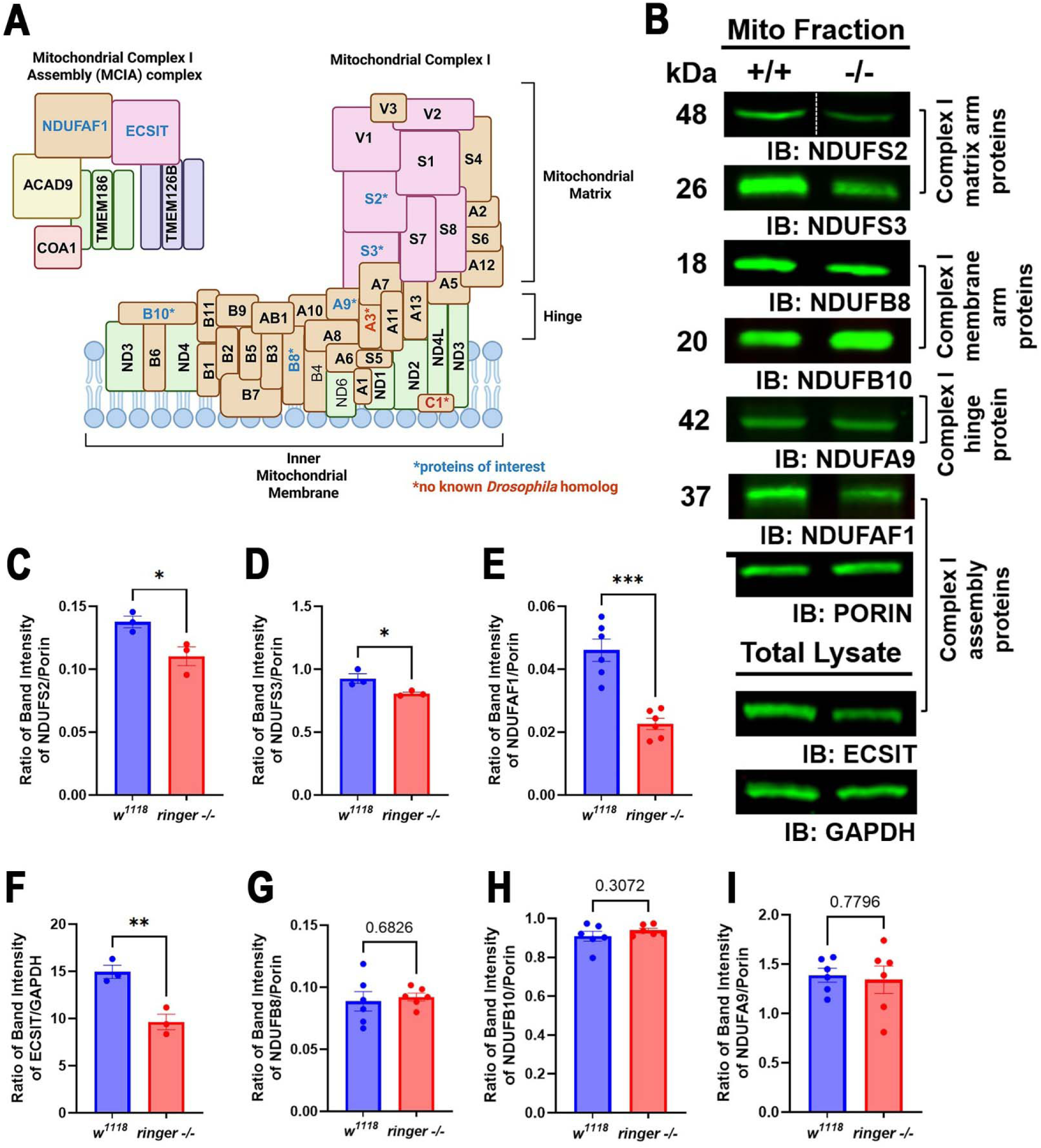
Loss of Ringer reduces the levels of mitochondrial complex I matrix subunits and assembly proteins. **A** Schematic of mitochondrial complex I and mitochondrial complex I assembly complex (MCIA) with individual subunits found in humans and flies. **B** Immunoblots of subcellular mitochondrial fractions from wild-type (+/+) and *ringer* mutant (-/-) fly heads probed with anti-NDUFS2 (48kDa), anti-NDUFS3 (26kDa), anti-NDUFB8 (18kDa), anti-NDUFB10 (20kDa), anti-NDUFA9 (42kDa), anti-NDUFAF1 (37kDa), and anti-Porin (30kDa) and total lysate fractions probed with anti-ECSIT (52kDa) and anti-GAPDH (37kDa). The white dotted line represents the space between nonconsecutive lanes of the same gel. Quantification of wild-type (lavender) and *ringer* mutant (red) complex I subunit and assembly protein levels **C** NDUFS2, t(4)=3.110, *p=0.0359, n=3, **D** NDUFS3, t(4)=3.076, *p=0.0371, n=3, **E** NDUFB8, t(10)=0.4211, p=0.6826, n=6, **F** NDUFB10, t(10)=1.076, p=0.3072, n=6, **G** NDUFA9, t(10)=0,2875, p=0.7796, n=6, **H** NDUFAF1, t(10)=5.848, ***p=0.0002, n=6, and **I** ECSIT, t(4)=4.910, **p=0.0080, n=3. Data are presented as mean ±SEM. Statistical significance was determined using a two-tailed unpaired t-test.

### Ringer loss alters CI protein density gradient profiles

Next, in order to determine: (i) if Ringer distributes in overlapping fractions with CI subunits, NDUFS2 and NDUFS3, and assembly proteins, NDUFAF1 and ECSIT in a buoyant density gradient and, (ii) if the distribution of these CI proteins is altered in *ringer* mutants, we performed Iodixanol/OptiPrep density gradient fractionation using ultracentrifugation from wild-type and *ringer* mutant brain extracts. Ringer was detected in immunoblots across the majority of fractions spanning the 10-40% density gradient in wild-type samples, indicating a broad distribution across the gradient (Fig. 3A,F), and was undetectable in *ringer* mutants. Ringer was detected in fractions containing NDUFS2 (Fig. 3B), NDUFS3 (Fig. 3C), ECSIT (Fig. 3D), and NDUFAF1 (Fig. 3E). In *ringer* mutants, NDUFS2 (Fig. 3G), NDUFS3 (Fig. 3H), ECSIT (Fig. 3I), and NDUFAF1 (Fig. 3J) exhibited subtle, yet consistent, shifts in their distributions. These data show that Ringer is enriched in fractions overlapping with CI matrix subunits and assembly proteins, and the loss of Ringer alters their distribution.

**Figure 3:**
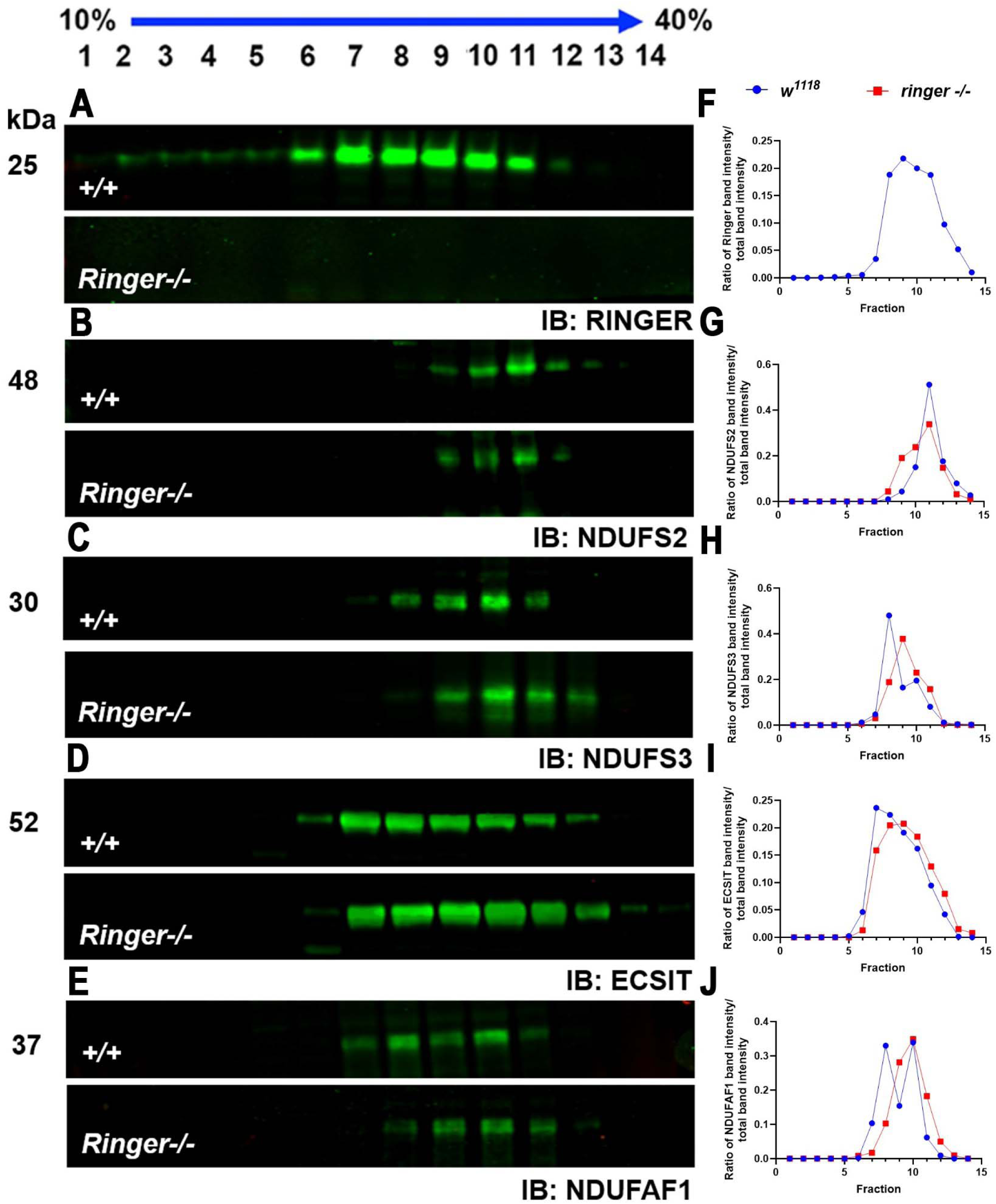
*ringer* mutants exhibit altered complex I subunit and assembly factor distribution across an iodixanol density gradient. Immunoblots of 10–40% OptiPrep density-gradient fractions probed with **A** anti-Ringer (25kDa), **B** anti-NDUFS2 (48kDa), **C** anti-NDUFS3 (26kDa), **D** anti-ECSIT (52kDa), and **E** anti-NDUFAF1 (37kDa) in wild-type (+/+) and *ringer* mutant (-/-) fly head lysates. Quantification of fraction intensity normalized to total intensity for **F** Ringer, **G** NDUFS2, **H** NDUFS3, **I** ECSIT, and **J** NDUFAF1, demonstrating profile shifts.

### *Ringer* mutants display reduced Complex I enzymatic activity

Since we observed a significant decrease in the protein levels of a subset of CI subunits in *ringer* mutants, we next investigated the consequences of Ringer loss on mitochondrial CI enzyme activity as a readout of CI function. CI catalyzes the oxidation of reduced nicotinamide adenine dinucleotide (NADH) to NAD+ and the reduction of ubiquinone (CoQ10) to ubiquinol (CoQH2).^32^ CI activity was quantified in mitochondrial lysates from adult *Drosophila* heads using a microplate-based assay in which absorbance was measured as a function of time (Fig. 4A). The slope of this graph was used to calculate the specific activity for each genotype with and without the addition of CI inhibitor, rotenone. *Ringer* mutants exhibited a significant reduction in CI-specific activity compared to wild-type controls (Fig. 4B), indicating a marked impairment in CI function. Consistent with assay specificity, rotenone treatment significantly decreased CI activity in both genotypes (Fig 4B), confirming that the measured rate reflected CI-dependent activity. These findings demonstrate that Ringer is vital for functional levels of CI enzymatic activity, and the loss of Ringer results in decreased NADH oxidation and ubiquinone reduction, potentially contributing to diminished ATP levels previously reported in *ringer* mutants.^6^

**Figure 4:**
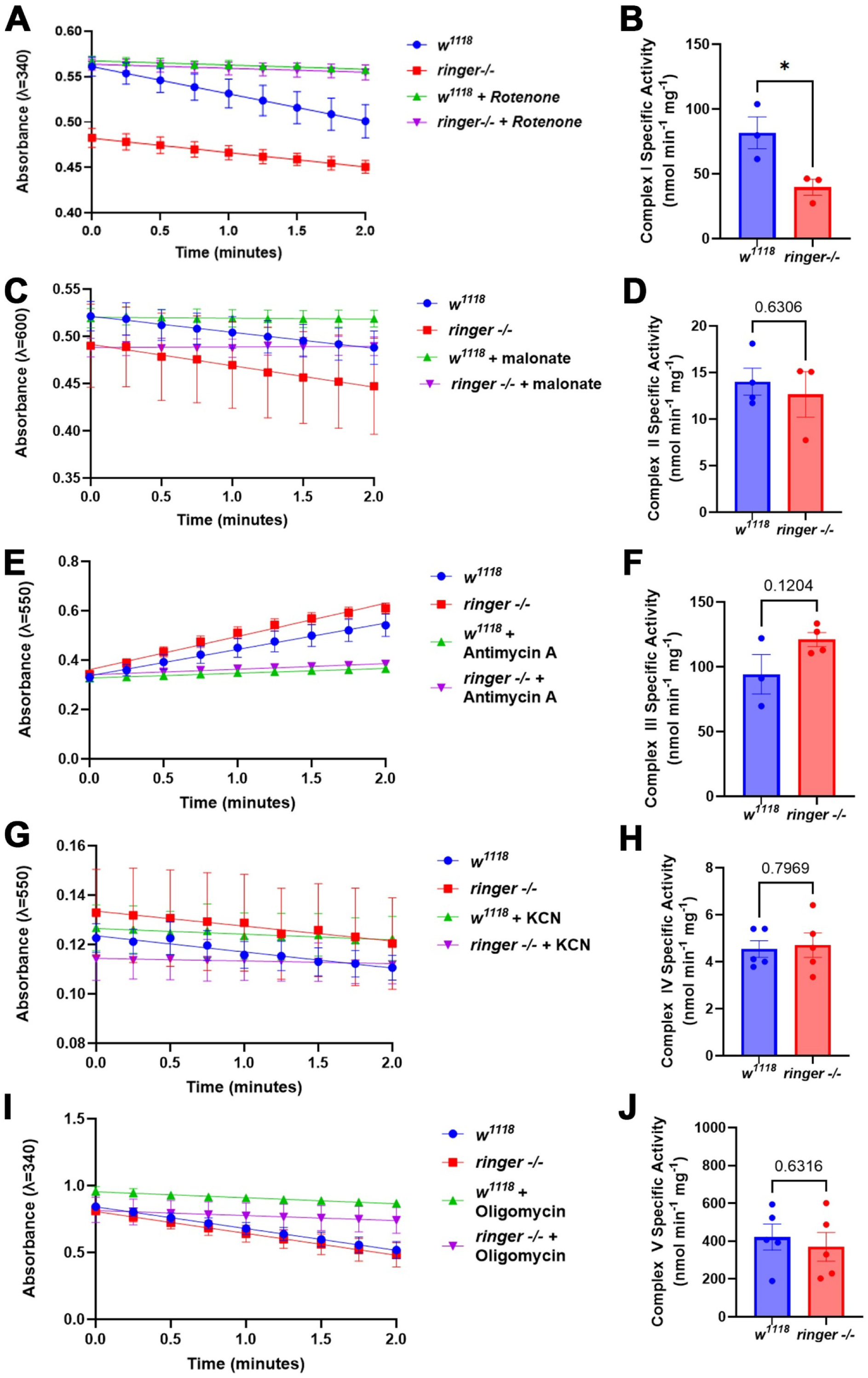
The loss of Ringer leads to impaired complex I enzymatic activity. **A** Plot of absorbance vs time graph depicting the decrease in absorbance for wild-type (blue), *ringer* mutant (red), wild-type + rotenone (green), and *ringer* mutant + rotenone (magenta) fly head lysate mitochondrial fractions. **B** Quantification of Complex I specific activity for wild-type control (n=3) and *ringer* mutant (n=3), t(4)=3.055, *p=0.0378. **C** Plot of absorbance vs time graph depicting the decrease in absorbance for wild-type (blue), *ringer* mutant (red), wild-type + Malonate (green), and *ringer* mutant + Malonate (magenta) fly head lysate mitochondrial fractions. **D** Quantification of Complex II specific activity for wild-type control (n=4) and *ringer* mutant (n=3), t(5)=0.5117, p=0.6306. **E** Plot of absorbance vs time graph depicting the decrease in absorbance for wild-type (blue), *ringer* mutant (red), wild-type + Antimycin A (green), and *ringer* mutant + Antimycin A (magenta) fly head lysate mitochondrial fractions. **F** Quantification of Complex III specific activity for wild-type control (n=3) and *ringer* mutant (n=4), t(5)=1.870, p=0.1204. **G** Plot of absorbance vs time graph depicting the decrease in absorbance for wild-type (blue), *ringer* mutant (red), wild-type + KCN (green), and *ringer* mutant + KCN (magenta) fly head lysate mitochondrial fractions. **H** Quantification of Complex IV specific activity for wild-type control (n=5) and *ringer* mutant (n=5), t(8)=0.2661, p=0.7969. **I** Plot of absorbance vs time graph depicting the decrease in absorbance for wild-type (blue), *ringer* mutant (red), wild-type + oligomycin (green), and *ringer* mutant + oligomycin (magenta) fly head lysate mitochondrial fractions. **J** Quantification of Complex V specific activity for wild-type control (n=5) and *ringer* mutant (n=5), t(8)=0.4984, p=0.6316. Data are presented as mean ±SEM. Statistical significance was determined using a two-tailed unpaired t-test.

Given that Ringer loss impairs CI enzymatic activity, we next evaluated whether this dysfunction extends to electron transport chain complexes II-V (Fig. 4C-J). We performed analogous enzyme activity assays for Complexes II–V using mitochondrial lysates from adult *Drosophila* heads. In contrast to the pronounced reduction in CI enzymatic activity, *ringer* mutants did not exhibit significant changes in the activities of CII (Fig. 4D), CIII (Fig. 4F), CIV (Fig. 4H), or CV (Fig. 4J) compared to wild-type controls following the subtraction of corresponding activities taken in the presence of complex II-V-specific inhibitors, malonate, antimycin A, potassium cyanide (KCN), and oligomycin, respectively. These findings indicate that the bioenergetic deficit in *ringer* mutants is selective for CI and may not reflect a global impairment of mitochondrial respiratory chain function.

### Resveratrol rescues mitochondrial deficits in *ringer* mutants

Given prior evidence that *ringer* mutants exhibit a significant increase in superoxide levels^6^, together with our current findings indicating an impaired CI function (Fig. 4), we hypothesize that in *ringer* mutants CI might be a major source of mitochondrial superoxide. We therefore next tested whether treatment with a ROS-scavenging compound, resveratrol,^33^ could mitigate these defects. Resveratrol directly interacts with CI, binding at or near the NADH-dehydrogenase domain to modulate electron transfer efficiency; this interaction stabilizes CI conformation, limits ROS-generating reverse electron transport, and supports improved catalytic activity under conditions of mitochondrial stress.^33^ Mitochondrial ROS levels were assessed using the MitoSOX assay in resveratrol-treated *ringer* mutants (Fig. 5A-C) compared to untreated *ringer* mutants that served as controls. Quantification of MitoSOX fluorescence intensity revealed a significant reduction in mitochondrial ROS in *ringer* mutants following resveratrol treatment (Fig. 5C), although levels remained elevated relative to wild-type animals (Supplemental Fig. 2A-C).

We previously reported that *ringer* mutants demonstrate reduced ATP levels.^6^ Therefore, we next tested whether resveratrol-treated *ringer* mutants would show an increase in ATP production compared to the untreated controls and found a significant increase, indicating a possible enhancement of oxidative phosphorylation (Fig. 5D). In parallel, CI enzymatic activity was also analyzed in untreated and resveratrol-treated *ringer* mutants. We found CI enzymatic activity to be significantly increased following resveratrol treatment in *ringer* mutants compared to untreated controls (Fig. 5E-F). However, lifespan analysis of resveratrol-treated *ringer* mutants compared to untreated ringer mutants (Supplemental Fig. 2E), showed no significant difference. As a control, we performed a lifespan analysis comparing untreated and resveratrol-treated wild-type flies (Supplemental Fig. 2D) and again saw no difference. In order to determine if resveratrol could promote neuronal survival in *ringer* mutants, neuronal counts performed by labeling with a pan-neuronal anti-Elav antibody in the resveratrol-treated and untreated *ringer* mutants (Fig. 5G,H) from two brain regions, the mushroom bodies (Fig. 5I) and the suboesophageal ganglia (Fig. 5J), also showed no significant difference. Together, these findings indicate that while resveratrol reduces mitochondrial ROS and improves CI-dependent bioenergetic function and increases ATP production in *ringer* mutants, it does not rescue neuronal degeneration or extend survival of *ringer* mutants.

**Figure 5:**
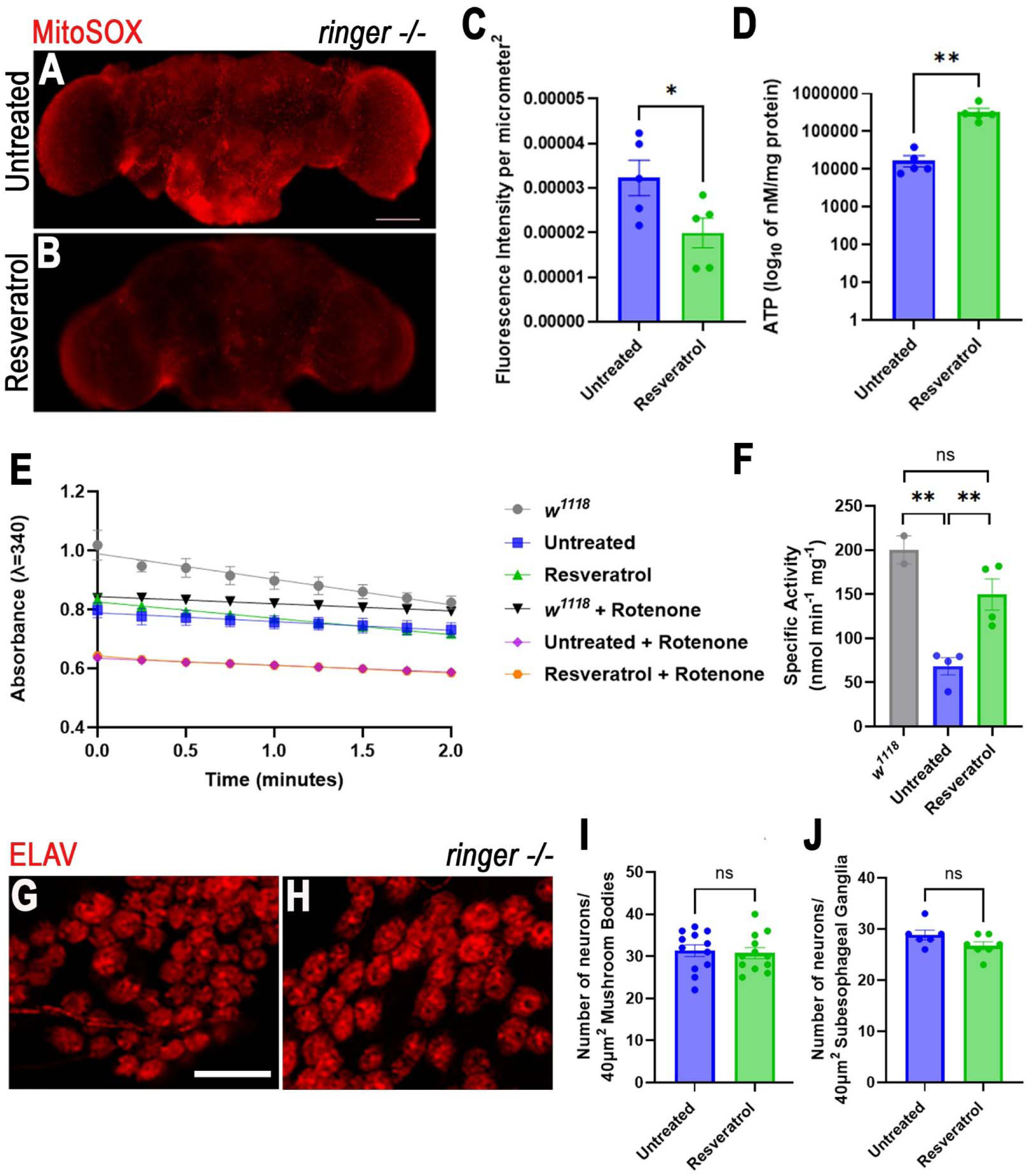
Resveratrol decrease ROS levels in ringer mutants compared to untreated controls. MitoSOX red fluorescence in live, unfixed whole-mount adult fly brains from **A** untreated and **B** resveratrol-treated *ringer* mutant flies. **C** Quantification of MitoSOX for *ringer* mutants untreated (n=5) and resveratrol (n=5). Data are presented as mean ±SEM. Statistical significance was determined using a two-tailed unpaired t-test, t(8)=2.368, *p=0.0454. Scale bar is 100µm. **D** Quantification of ATP values for *ringer* mutants treated with resveratrol compared with untreated controls plotted as the log₁₀-transformed values of ATP concentration (log₁₀ nM/mg of protein). Data are presented as mean ±SEM. Statistical significance was determined using a two-tailed unpaired t-test, t(8)=3.921, **p=0.0044. **E** Plot of absorbance vs time graph depicting the decrease in absorbance for wild-type (grey), *ringer* mutant (blue), resveratrol-treated *ringer* mutants (green), wild-type + rotenone (black), *ringer* mutant + rotenone (magenta), and resveratrol-treated *ringer* mutants (orange) fly head lysate mitochondrial fractions. **F** Quantification of complex I specific activity for wild-type flies (n=2), *ringer* mutants treated with resveratrol (n=4) and *ringer* mutant untreated controls (n=4). Data are presented as mean ±SEM. Statistical significance was determined using a two-tailed unpaired t-test. wild-type vs untreated *ringer* mutants t(4)=7.549, **p=0.0016, untreated *ringer* mutants vs resveratrol-treated mutants t(6)=4.030, **p=0.0069, and wild-type vs resveratrol-treated *ringer* mutants t(4)=1.792, p=0.1476. **G** ELAV fluorescence in the mushroom bodies of adult fly brains from **G** untreated and **H** resveratrol-treated *ringer* mutant flies. Quantification of neurons per 40µm^2^ in the **I** mushroom bodies, t(22)=0.3062, p=0.7623, and **J** suboesophageal ganglia, t(11)=1.747, p=0.1084, in adult fly brains. Data are presented as mean ±SEM. Statistical significance was determined using a two-tailed unpaired t-test.

### TPPP Localization in BA40 of Control and PD Human Cortex

Given that mitochondrial dysfunction is found in many neurodegenerative diseases, including PD, and human TPPP is implicated in PD, we next examined whether TPPP shows altered localization in human PD brain tissue.^34^ Based on our tissue availability from the parietal cortex BA40 region, we performed immunostaining on BA40 sections from age-matched healthy control and PD male subjects, which revealed marked differences in TPPP localization (Fig. 6A-B”). TPPP is enriched in the oligodendrocytes in control BA-40 tissue (white arrows, Fig. 6A”, A”), as revealed by co-labeling with Connexin-32, an oligodendrocyte-associated protein (white arrows, Fig. 6A, A”). In the PD BA40, the subcellular localization of TPPP is altered when compared with controls and is accompanied with aggregate formation (Fig. 6B’, B”, D). These findings indicate a clear dysregulation of TPPP in the PD BA40 region of the parietal cortex.

**Figure 6:**
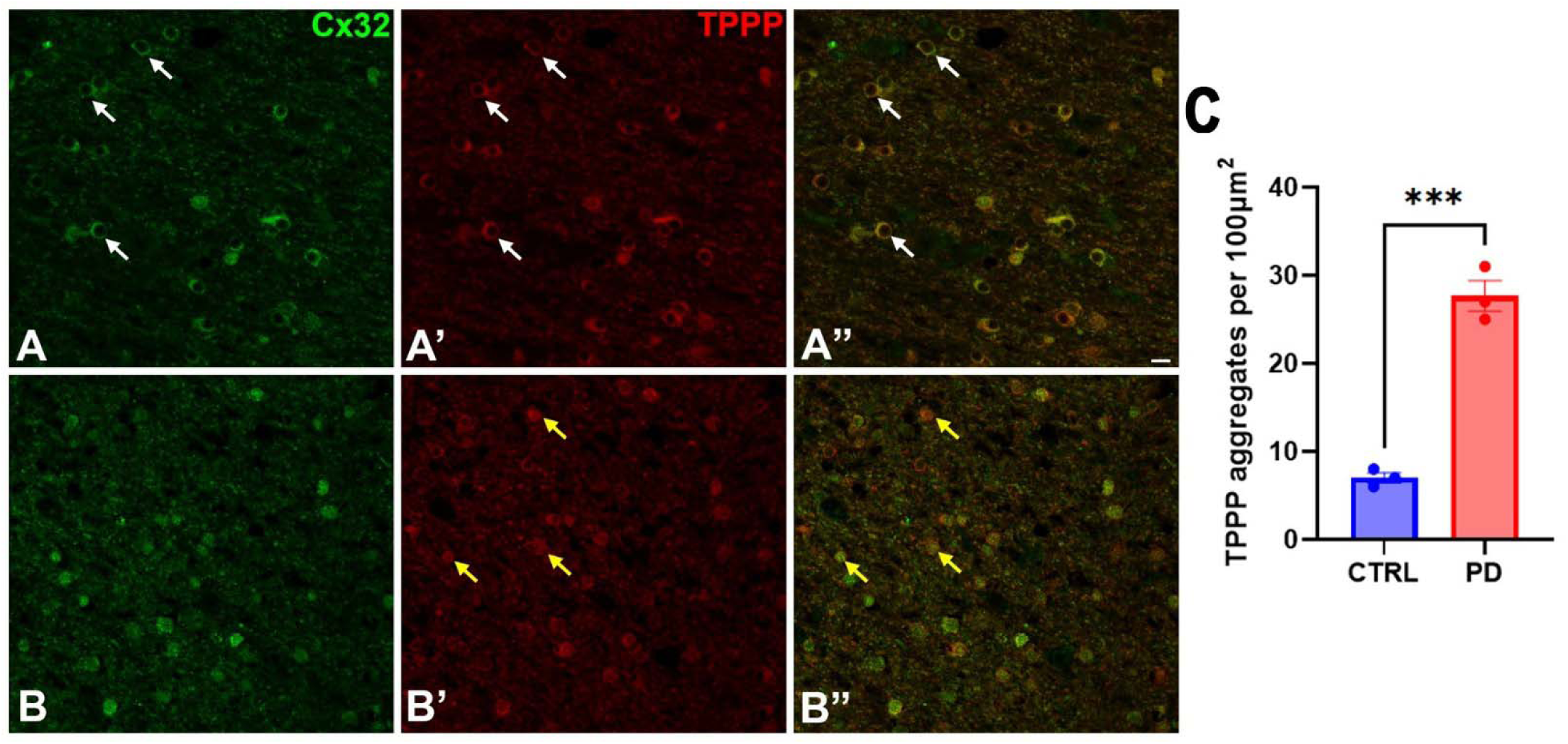
TPPP aggregates in the BA40 region of the PD brain compared to healthy controls. Confocal images of BA40 sections from age-matched **A-A”** healthy controls n=3 and **B-B”** PD n=3 male subjects co-labeled with Cx32 (green) and TPPP (red). **C** Quantification of the number of TPPP aggregates per 100µm^2^. Data are presented as mean ±SEM. Statistical significance was determined using a two-tailed unpaired t-test, t(4)=11.14, ***p=0.0004.

### Specific CI proteins are reduced in the PD parietal cortex

In order to validate the findings from our *Drosophila ringer* mutant model and examine its relevance to PD, we next assessed whether the levels of the same CI subunits analyzed in the *ringer* mutants are also altered in human PD brain tissue from the BA40 region of the parietal cortex. Total lysate brain homogenates from postmortem male PD patients and respective age- and sex-matched controls were processed for immunoblotting (Table 1, Fig. 7A). Anti-GAPDH was used as a loading control. We quantified the ratio of the band intensities of NDUFS2, NDUFAF1, NDUFS3, NDUFB8, NDUFB10, NDUFA9, and ECSIT with GAPDH (Fig. 7A). In PD samples, core CI matrix-arm subunit NDUFS2 (Fig. 7B) as well as the assembly factor NDUFAF1 (Fig. 7C) were significantly reduced compared to controls. In contrast, matrix protein NDUFS3 (Fig. 7D), membrane-arm subunits NDUFB8 (Fig. 7E) and NDUFB10 (Fig. 7F), hinge subunit NDUFA9 (Fig.7G), and assembly factor ECSIT (Fig. 7H) did not show significant differences between the control and PD groups. While these findings show a correlation between human PD and our *Drosophila* model, the fly model does not fully recapitulate the findings from the human PD tissues.

**Figure 7:**
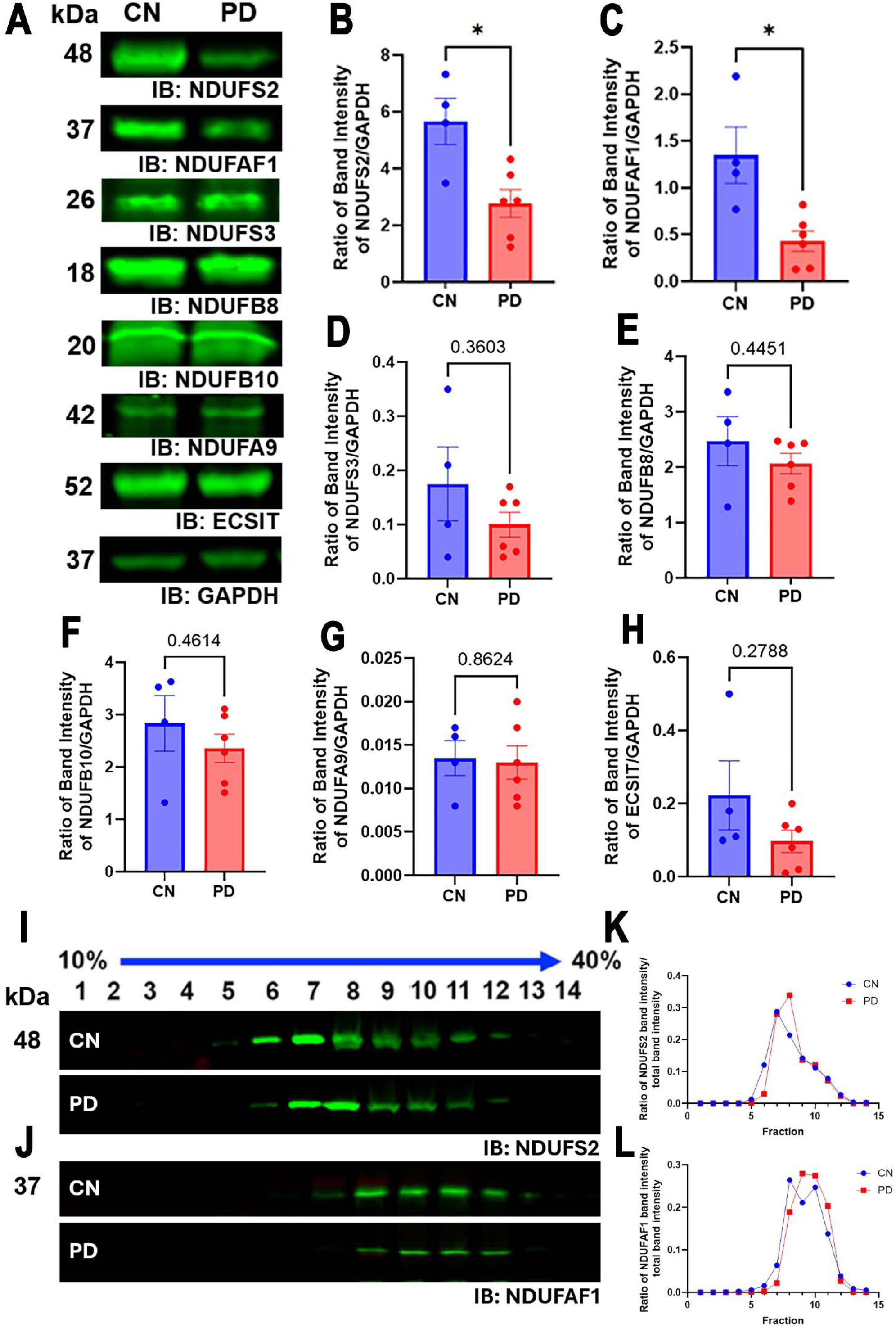
Complex I matrix and assembly proteins are reduced in Brodmann Area 40 of Parkinson’s disease brains and show altered distribution across iodixanol density gradient fractions. **A** Immunoblots of total lysate fractions from human male control and PD postmortem brain tissue lysates from the BA40 region of the parietal cortex probed with anti-NDUFS2 (48kDa), anti-NDUFAF1 (37kDa), anti-NDUFS3 (26kDa), anti-NDUFB8 (18kDa), anti-NDUFB10 (20kDa), anti-ECSIT (52kDa), and anti-GAPDH (37kDa). Quantification of control (CN, blue) and PD (red) complex I subunit and assembly protein levels **B** NDUFS2, t(5.212)=3.062, *p=0.0359; **C** NDUFAF1, t(3.811)=2.862, *p=0.0485, **D** NDUFS3, t(3.691)=1.043, **E** NDUFB8, t(4.080)=0.8445, **F** NDUFB10, t(4.541)=0.8040, **G** NDUFA9, t(7.283)=0.1796, and **H** ECSIT, t(3.625)=1.272. . For control samples, n=4, and for PD samples, n=6. Data are presented as mean ±SEM. Statistical significance was determined using a two-tailed unpaired t-test with Welch’s correction. Immunoblots of 10–40% OptiPrep density-gradient fractions probed with **I** anti-NDUFS2 (48kDa) and **J** anti-NDUFAF1 (37kDa) in human male control and PD postmortem brain tissue lysates from the BA40 region of the parietal cortex. Quantification of fraction intensity normalized to total intensity for **K** NDUFS2and **L** NDUFAF1, demonstrating profile shifts.

**Table 1:** Male subject demographics and neuropathological profiles for control and Parkinson’s disease patient cohorts. Parietal cortex Brodmann area 40 (BA40) from frozen human brain tissue samples were obtained from the NIH NeuroBioBank. Male subjects with Parkinson’s disease (PD, n=6) and age-matched controls (CN, n=4) were selected for comparison via molecular approaches, with case information summarized above. CN and PD male subjects’ ages are reported in years, and the postmortem interval (PMI) is reported in hours. Neuropathological assessments include Braak/Del Tredici staging (Stage 1-6) and Lewy Body disease (LBD) classifications representative of α-synuclein pathology. Higher Braak/Del Tredici stages indicate more extensive progression of PD-related pathology in the parietal cortex, while limbic and diffuse LBD classifications describe the anatomical distribution of Lewy Body pathology.

| Diagnosis | Subject | Age<br>(years) | PMI<br>(hours) | Neuropathology |
| --- | --- | --- | --- | --- |
| Unaffected<br>Control (CN) | CN-1 | 84 | 28.8 | None |
|  | CN-2 | 83 | 13.0 | None |
|  | CN-3 | 80 | 18.3 | None |
|  | CN-4 | 81 | 13.7 | None |
| Parkinson's<br>Disease (PD) | PD-1 | 83 | 15.0 | Braak stage 4 |
|  | PD-2 | 84 | 13.2 | Diffuse LBD, limbic stage |
|  | PD-3 | 85 | 22.1 | Braak/Del <u>Tredici</u> stage 5 |
|  | PD-4 | 81 | 19.2 | Braak/Del <u>Tredici</u> stage 4 |
|  | PD-5 | 80 | 12.8 | Early diffuse LDB |
|  | PD-6 | 80 | 13.4 | Braak/Del <u>Tredici</u> stage 4 |

To evaluate the distribution of NDUFS2 and NDUFAF1 in control and PD human brain tissues, we performed similar Iodixanol/OptiPrep density-gradient fractionation on BA40 samples. In control tissue, CI matrix subunit NDUFS2 (Fig. 7I,K) and the assembly factors NDUFAF1 (Fig. 7J,L), localized to defined fractions characteristic of CI-associated protein pools consistent with fly analyses (Fig. 3). In PD samples, these CI proteins were detected mostly in overlapping fractions as in the control but exhibited an overall reduced distribution across the density-gradient fractions (Fig. 7I-L). Together, these results demonstrate that select CI components disrupted in *ringer* mutants show parallel reductions in human PD BA40 tissue, underscoring a conserved vulnerability of CI matrix-arm subunits and assembly factors in pathological conditions. Although correlative, the convergence of biochemical deficits and distributional changes of proteins in both *Drosophila* and humans supports the relevance of the *ringer* model to further investigate PD-associated mitochondrial dysfunction.

## DISCUSSION

Our findings identify Ringer as a previously unrecognized regulator of mitochondrial CI assembly and function, revealing a role for TPPP homologs in maintaining mitochondrial bioenergetics. We show that Ringer is associated with both the cytoplasm and the mitochondria equally and is specifically enriched in the mitochondrial matrix. In addition to the marked mitochondrial ultrastructural abnormalities shown previously,^6^ our findings reveal that loss of Ringer causes bioenergetic deficits, including reduced levels of multiple CI matrix subunits and assembly factors, highlighting deficits in specific CI subunits and associated proteins with Ringer loss. These molecular defects are accompanied by a significant reduction in CI enzymatic activity without impacting the enzymatic activities of the other complexes in the ETC, establishing a clear functional consequence of Ringer function in the mitochondrial CI. Some of the changes, but not all, to specific CI subunit levels seen in the *ringer* mutants are recapitulated in human PD, together with altered TPPP localization in the BA40 region of the brain.

The central insight from our work is that Ringer regulates specific CI subunit abundance and enzymatic activity within the mitochondrial matrix, as evidenced by the reduction in levels of specific core CI subunits and assembly factors essential for CI function. Among these, ECSIT and NDUFAF1 form a critical assembly module required for the stabilization and maturation of the matrix and membrane arms of CI.^35^ Mutations or depletion of either protein cause severe mitochondrial encephalopathies or CI deficiency disorders.^31^ Likewise, NDUFS2 and NDUFS3 are essential catalytic core subunits that form part of the N module responsible for NADH oxidation, and pathogenic variants in these genes lead to Leigh syndrome, progressive neurodegeneration, and/or profound respiratory chain impairments.^36^ The reduction of these factors in *ringer* mutants reflects the disruption of highly conserved steps in CI biogenesis and catalytic function.

The specific vulnerabilities associated with each of these components further reinforce the mechanistic significance of their loss. Downregulation of NDUFS2 in *Drosophila* activates TOR signaling, accelerating aging, promoting neurodegeneration, and diminishing immune competence,^37^ suggesting that reduced NDUFS2 levels in *ringer* mutants may trigger maladaptive stress-response pathways that exacerbate mitochondrial dysfunction. NDUFS3 loss is similarly detrimental, as mutations in NDUFS3 can cause encephalopathies.^38,39^ NDUFS3 ablation in mouse models results in early myopathy and mortality,^40^ and partial NDUFS3 depletion destabilizes the entire CI holoenzyme,^40^ indicating that even modest reductions in NDUFS3, such as those observed in *ringer* mutants, could compromise CI holoenzyme integrity. ECSIT loss has been linked to impaired mitochondrial ROS signaling, defective mitophagy, loss of the CI holoenzyme,^41^ and a paradoxical increase in PARKIN levels without corresponding increases in mitophagy,^42^ a phenotype highly relevant to PD given PARKIN’s propensity to aggregate in disease.^43^ The reduction of ECSIT in *ringer* mutants, therefore, suggests a possible convergence of TPPP dysfunction, impaired CI assembly, and dysregulated mitochondrial quality control.

The localization of Ringer to the mitochondrial matrix provides a spatial basis for these effects and raises the possibility that TPPP homologs act as scaffolding or chaperone-like elements that stabilize early assembly intermediates. This interpretation aligns with emerging evidence that TPPP1 influences protein aggregation and organelle homeostasis, suggesting that TPPP-dependent pathways may intersect with mitochondrial proteostasis in ways that have not been previously demonstrated.^44^ Moreover, because Ringer is selectively enriched within the mitochondrial matrix, it is possible that this spatial proximity underlies why matrix-localized CI components are disproportionately affected by Ringer loss while the membrane arm and hinge subunits analyzed remain unaffected.

In *ringer* mutants, CI matrix-arm subunits and assembly factors also exhibited modest shifts across the Iodixanol density gradient, indicating that CI organization might be altered under conditions of Ringer loss which needs to be further examined. In healthy mitochondria, CI components distribute across discrete fractions.^45^ The rightward shifts we observe likely reflect the accumulation of incompletely or inappropriately assembled intermediates, as well as the enhanced burden of damaged mitochondria.^46^ Conversely, the leftward shifts detected for a subset of CI proteins might suggest the presence of smaller, destabilized subcomplexes or prematurely disassembled intermediates, indicating that Ringer loss may perturb both the formation and maintenance of CI structures.^47^

The ability of resveratrol to ameliorate ROS levels and improve CI activity and ATP levels in *ringer* mutants highlights the therapeutic potential of targeting oxidative stress in the context of TPPP dysfunction. Resveratrol has been shown to activate SIRT1,^48^ stimulate PGC-1α–dependent mitochondrial biogenesis,^49^ enhance antioxidant defenses,^50^ and promote mitophagy and mitochondrial turnover in multiple model systems.^51,52^ It can also stabilize CI activity under conditions of oxidative or metabolic stress and has been reported to preserve NAD⁺ homeostasis, a key determinant of CI function.^53^ Together, these mechanisms raise the possibility that resveratrol counteracts the consequences of Ringer loss by buffering ROS-driven mitochondrial damage. Its efficacy in reducing ROS levels in *ringer* mutants suggests that the mitochondrial defects arising from Ringer loss are at least partially reversible and responsive to interventions that bolster mitochondrial resilience.

The inability of resveratrol to rescue neurodegeneration or survival in *ringer* mutants indicates that Ringer dysfunction triggers additional pathogenic processes beyond elevated ROS, consistent with the multifunctional nature of Ringer. As such, rescues to neurodegeneration and survival will likely require combination therapies that target multiple mechanisms rather than antioxidant treatment alone, similar to the multifactorial nature of PD pathogenesis. Notably, mitochondrial impairments seen in *ringer* mutants parallel key pathological features observed in PD, and our analyses of human postmortem tissue from parietal cortex BA40 region reveal similar decreases in CI proteins NDUFS2 and NDUFAF1, underscoring the translational relevance of our model.

Despite the many strengths of the studies, there are some limitations that should be acknowledged. First, our analyses focused primarily on CI subunits and assembly factors and therefore do not capture the full extent of mitochondrial dysfunction that may arise from Ringer loss. Second, while Optiprep experiments indicate the likelihood of Ringer association with specific CI subunits, it does not confirm protein-protein interactions. Additional experiments, including co-immunoprecipitations, which will be part of our future investigations are needed to provide direct evidence of Ringer-CI interactions, as well as elucidate specific mechanistic roles that Ringer might play in CI assembly and function. Additionally, while Optiprep gradient experiments can detect distribution changes of proteins across fractions, which is informative; however, it does not provide sufficient resolution to clearly separate cellular compartments and sub-compartments. Therefore, they do not support definitive conclusions regarding a compartment-specific shift in protein distributions. Third, although these *Drosophila*-human parallels strengthen the relevance of the *ringer* mutant, we note that the fly model used in this study is a knockout model and therefore may exaggerate CI-related phenotypes relative to human PD, where TPPP is not deleted. Fourth, the extent of changes in CI proteins may be brain region-specific which may also contribute to the incomplete recapitulation of the molecular changes observed in human PD tissues, as the Drosophila model may not capture the regional heterogeneity observed in the human brain. Finally, due to the limitations in tissue availability, the human biochemical data in the current study were limited to total homogenates as opposed to mitochondrial fractions as shown in the *Drosophila*, as well as only male subjects and the BA40 region of the parietal cortex. Although the BA40 of the parietal cortex is implicated in PD pathology, it is not the primary site of dopaminergic neurodegeneration. Future analyses of mitochondrial fractions in more vulnerable regions such as the substantia nigra and in female subjects will be essential for defining the extent of overlap between the *Drosophila* and human PD mitochondrial deficits as well as sex-specific differences to TPPP dysfunction in PD.

Together, these approaches as part of our future investigations will clarify whether Ringer/TPPP regulates mitochondrial function through CI-specific mechanisms or through a multi-complex mechanism and metabolic pathways, thereby providing deeper insight into how TPPP dysfunction may drive mitochondrial impairment in neurodegeneration and PD. This model provides an entry point and enables direct exploration of how conserved TPPP-dependent mechanisms intersect with mitochondrial health, providing an effective *in vivo* model to elucidate the mechanistic pathways through which TPPP dysfunction drives mitochondrial deficits in synucleinopathies.

## METHODS

### Drosophila stocks

The *Drosophila* stocks utilized in this study include *w^1118^* (wild type control), obtained from the Bloomington Drosophila Stock Center (stock #3605), and *ringer^915.3^* All *Drosophila* stocks were maintained at 25 °C under controlled environmental conditions, including 50% relative humidity and a 12-hour light/dark cycle. Male and female flies used for all experiments were age matched. Day-0 adults were collected upon eclosion, placed into fresh food vials, and maintained under standard husbandry conditions, with transfers into fresh vials performed twice weekly to preserve cohort health.

### Subcellular and sub-mitochondrial fractionations

Subcellular fractionations were performed using adult *Drosophila* heads to isolate total lysates, mitochondrial fractions, and cytoplasmic fractions from wild-type and *ringer* mutant flies as previously described.^4^ Briefly, heads were homogenized in mitochondrial isolation buffer (mito buffer; 250mM sucrose, 10mM Tris-HCl, 1mM EGTA, pH 7.5), debris was removed by low-speed centrifugation (800g for 5 minutes), and the resulting supernatant was centrifuged at 10,000g for 20 minutes to obtain the crude mitochondrial pellet and post-mitochondrial (cytoplasmic) supernatant. Three to six biological samples per genotype were prepared for all fractionation experiments. Mitochondrial fractions were resuspended in mitochondrial isolation buffer, quantified using Micro BCA kit (ThermoFisher, #23235), resolved on 10% or 12% SDS–PAGE gels, subjected to immunoblotting, and probed with antibodies against CI subunits and assembly factors and loading controls.

For mitochondrial matrix and membrane separation, mitochondria isolated from wild-type adult brain lysates were further fractionated using a sodium carbonate extraction method. The mitochondrial pellet was resuspended in 50 µL of 0.1 M sodium carbonate, and 20 µL of this suspension was retained as the total mitochondrial fraction. The remaining 30 µL was incubated on ice for 30 min to promote osmotic disruption and protein solubilization. Samples were then ultracentrifuged at 42,000 rpm for 40 min at 4 °C. The supernatant, representing the matrix fraction, was transferred to a new tube, while the pellet containing the membrane fraction was resuspended in 30 µL of 0.1 M sodium carbonate. Total mitochondrial, matrix, and membrane fractions were resolved on 10% SDS–PAGE gels and analyzed by immunoblotting.

### OptiPrep (iodixanol) density gradient fractionation

OptiPrep (iodixanol) density-gradient fractionation was performed to resolve proteins based on density.^47^ Adult fly heads or human brain tissues were homogenized in 1% Triton-X lysis buffer with protease and phosphatase inhibitors and cleared by low-speed centrifugation to remove debris. The supernatant was layered onto a 10–40% step OptiPrep gradient prepared in lysis buffer and ultracentrifuged at 25,000 rpm for 4 hours to allow separation of protein complexes across the gradient. Following centrifugation, 14 fractions were collected sequentially from top to bottom in equal volumes. Each fraction was mixed with SDS sample buffer, resolved on 10% or 12% SDS-PAGE gels, and analyzed by immunoblotting to determine the distribution of CI matrix subunits and assembly factors across density layers.

### Immunoblotting

Adult fly heads and human postmortem tissue were processed for immunoblotting following established protocols. Equal amounts of protein from *Drosophila* head homogenates were resolved by SDS-PAGE and probed with the appropriate primary antibodies, with 3–6 biological replicates per experiment.^6^ Similarly, frozen human tissue total lysate samples were generated by homogenizing tissue in 1% Triton-X lysis buffer at a 1:10 (w/v) ratio, quantified by BCA assay (ThermoFisher, #23227), and equal protein amounts were subsequently resolved by SDS-PAGE and immunoblotted with the appropriate primary antibodies.^54^ Primary antibodies used for immunoblotting were anti-Ringer (1:10,000), anti-GAPDH (1:5000, Invitrogen, MA5-15,738-BTIN), anti-Porin (1:2000, Abcam, ab14734), anti-PDH (1:500, 126224), anti-NDUFS2 (1:4000, Abcam, 192022), anti-NDUFS3(1:2000, Abcam, 177471), anti-NDUFB8 (1:2000, Abcam, 192878), anti-NDUFB10 (1:5000, Abcam, 196019), NDUFA9(1:1000, Abcam, 14713), anti-ECSIT(1:1000, Abcam, 21288), and anti-NDUFAF1(1:5000, Abcam, 79826).

### ETC complex enzymatic activity assay

CI enzymatic activity was measured in adult wild-type and *ringer* mutant fly heads using a previously standardized microplate-based assay with minor modifications.^55^ For all enzymatic activity assays, samples were prepared from adult wild-type and ringer mutant fly heads according to the mitochondrial fractionation protocol above. Mitochondria were isolated and resuspended in isotonic buffer (10 mM Tris-HCl, pH 7.6) at a final concentration of 1 µg/µL and subjected to three liquid-nitrogen freeze–thaw cycles. To measure complex I enzymatic activity, 10 µL of mitochondrial lysate was added to 186 µL of reaction mix (50 mM potassium phosphate buffer pH 7.5, 3mg/mL BSA, 300µM KCN, 100µM NADH) in each well of a 96-well plate in the absence and presence of Complex I inhibitor rotenone. Baseline absorbance at 340 nm was recorded at 30 °C before initiating the reaction with 4 µL of 4 mM CoQ₁. Absorbance at 340 nm was then monitored for an additional 2 min at 30 °C. To measure complex II enzymatic activity, 10 µL of mitochondrial lysate was added to 186 µL of reaction mix (25 mM potassium phosphate buffer pH 7.5, 1mg/mL BSA, 300µM KCN, 20mM succinate, 75µM DCPIP) in each well of a 96-well plate in the absence and presence of Complex II inhibitor malonate. Following a 10-minute incubation step at room temperature, baseline absorbance at 600 nm was recorded at 30 °C before initiating the reaction with 2 µL of 4 mM CoQ₁. Absorbance at 600 nm was then monitored for an additional 2 min at 30 °C. To measure complex III enzymatic activity, 10 µL of mitochondrial lysate was added to 186 µL of reaction mix (25 mM potassium phosphate buffer pH 7.5, 100µM EDTA pH 7.5, 500µM KCN, 0.025% Tween-20, 75µM oxidized cytochrome c) in each well of a 96-well plate in the absence and presence of Complex III inhibitor antimycin A. Baseline absorbance at 550 nm was recorded at 30 °C before initiating the reaction with 2 μL of 10 mM of decylubiquinol. Absorbance at 550 nm was then monitored for an additional 2 min at 30 °C. To measure complex IV enzymatic activity, 10 µL of mitochondrial lysate was added to 188 µL of reaction mix (50mM potassium phosphate buffer pH 7.0, 75mM reduced cytochrome c) in each well of a 96-well plate to start the reaction. This was performed in the absence and presence of Complex IV inhibitor 500µM KCN. Absorbance at 550 nm was monitored for 2 min at 30 °C. To measure complex V enzymatic activity, 10 µL of mitochondrial lysate was added to 170 µL of reaction mix (50uM HEPES buffer pH 8.0, 0.3mM NADH, 2.5mM phosphoenolpyruvate, 50mg/mL pyruvate kinase,50mg/mL lactate dehydrogenase, 2mg/mL antimycin A) in each well of a 96-well plate in the absence and presence of Complex V inhibitor oligomycin. Following a 5-minute incubation at room temperature, baseline absorbance at 340 nm was recorded at 30 °C before initiating the reaction with 20 µL of 25 mM ATP. Absorbance at 340 nm was then monitored for an additional 2 min at 30 °C. Specific activity was calculated from the linear slope of absorbance over time using linear regression analysis.

### Pharmacological treatments

Drug treatments were performed using resveratrol^56^ with minor adaptations to previously described methods. Freshly eclosed wild-type and *ringer* mutant flies (Day 0) were transferred to food vials containing filter paper saturated with resveratrol or the vehicle (50/50 ethanol and 10% sucrose in water). Resveratrol was dissolved in 100% ethanol and diluted to 50% ethanol with a 10% sucrose solution and applied to filter paper at a final concentration of 2.5 mM. The filter paper was then inserted into a standard food vial. Flies were transferred to freshly prepared vials every third day for the duration of the assays.

### MitoSOX ROS assay

Mitochondrial ROS were measured using the MitoSOX Red mitochondrial superoxide indicator as described previously.^6^ Briefly, adult fly brains were dissected in ice-cold 1X PBS and immediately transferred to staining solution containing 5 µM MitoSOX Red prepared fresh in 1X PBS. Tissues were incubated for 15 minutes at 37°C in the dark to prevent photo-oxidation. Following incubation, brains were rinsed in 1X PBS and mounted on Vectasheild antifade mounting medium (Vector Laboratories, H-1000-10) on slides. All samples were imaged on a Keyence BZ-X710 fluorescence imaging system using identical laser power, exposure time, and detector gain settings across genotypes and drug treatment groups to ensure consistency. Fluorescence intensity was quantified in ImageJ by calculating the mean intensity value per micron^2^ across each brain. These values were treated as independent biological replicates. A total of n=5-6 brains per genotype were analyzed for all MitoSOX measurements.

### ATP assay

ATP levels were measured using the Promega Enliten kit (Promega, FF2000) based on previously established protocols.^57^ Briefly, six age-matched control or resveratrol-treated flies were homogenized in 50 μl of extraction buffer (6 M guanidine-HCl, 100 mM Tris and 4 mM EDTA, pH 7.75). Samples were frozen in liquid nitrogen, thawed, spun at 14,000 rpm for 15 minutes, and the supernatant collected. Following a BCA assay to determine protein concentration, samples were diluted (1:2000) with dilution buffer (100 mM Tris and 4 mM EDTA, pH 7.75) and mixed with Enliten luminescent solution. Luminescence was measured on the SpectraMax iD3 plate reader. ATP molarity was determined by plotting and standard curve and normalized to the concentration of protein per sample.

### Lifespan assay

Lifespan analysis was conducted based on previously established methods.^6^ Briefly, 50 flies (25 males and 25 females) were used for the lifespan assay. Wild-type and *ringer* mutant fly survival was measured in food vials containing filter paper soaked in the vehicle (equal volume EtOH and 10% sucrose), or 2.5mM Resveratrol ^26,58^. Flies were transferred to fresh food every third day and maintained at 25 °C throughout the assay. Survival curves were generated using Kaplan-Meier analysis in GraphPad Prism.

### Fly Brain Immunostaining

Immunostaining of whole-mount adult fly brains was performed based on previous protocols.^6^ Briefly, adult fly brains were dissected in ice-cold PBS and followed by a 20-minute fixation in 4% paraformaldehyde (PFA) prepared in PBS for 20 min, followed by staining with primary and secondary antibodies. The primary antibody used was anti-Elav (1:500, DSHB, 9F8A9), and the secondary antibody conjugated to Alexa 647 was used at a 1:400 dilution. Whole brains were then mounted with Vectasheild antifade mounting medium (Vector Laboratories, H-1000-10) on slides, and confocal images were acquired. To quantify neuronal density, Elav-positive nuclei were counted within a 40 × 40 μm^2^.

### Human Brain Immunostaining

Formalin-fixed, paraffin-embedded human BA40 brain sections were deparaffinized in xylene (10 min, then 5 min) and rehydrated through a graded ethanol series (100% for 3 min x2, 95% for 2 min, 80% for 2 min) followed by two rinses in deionized water. Slides were then exposed to photo-irradiation in citrate buffer for 2 h at 4 °C and subjected to heat-mediated antigen retrieval in 1x citrate buffer using a pre-heated steamer (95–100 °C) for 20 min. After cooling for 5 min, sections were washed in PBS, then PBT before blocking in 2% BSA in PBT for 1 h. Primary antibodies diluted in 2% BSA/PBT were applied overnight at 4 °C. The following morning, slides were washed in PBS and PBT and incubated with Alexa Fluor–conjugated secondary antibodies (1:400) for 2 h at room temperature in the dark. After final washes in PBT and PBS, excess buffer was removed, and tissues were cover-slipped with mounting medium to preserve fluorescence.

### Statistical Analysis

Western blot analyses were performed on mitochondrial fractions and total lysate samples to quantify steady-state levels of CI subunits and assembly factors. Band intensities were measured from fluorescence images using LiCOR software. For each blot, signal intensities were normalized to loading controls (Porin for mitochondrial fractions or GAPDH for total lysates). All genotypes within a given experiment were processed, transferred, and imaged in parallel under identical conditions. A total of n=3-6 biological replicates per genotype were analyzed for both human and *Drosophila* studies.

CI enzymatic activity was quantified using a microplate-based absorbance assay. Mitochondrial fractions were isolated from adult fly heads, and C1 enzymatic activity was measured by monitoring the decrease in absorbance at 340 nm. Activity values were calculated using the linear regression slope calculated in Excel from the change in absorbance over two minutes. n = 3-4 biological replicates per genotype were included for each assay.

Mitochondrial superoxide levels were quantified using the MitoSOX Red assay. Samples from all genotypes were imaged using identical Keyence acquisition parameters to ensure comparability across groups. Fluorescence intensity was measured in ImageJ from the whole fly brain. Mean fluorescence values were calculated for each brain and normalized over the total area. A total of n=5-6 brains per genotype were analyzed for all MitoSOX quantifications.

Survival was recorded for a span of four weeks. Survival curves were generated in GraphPad Prism using the Kaplan-Meier method, and statistical significance was assessed using the log-rank (Mantel–Cox) test. Median and maximum lifespan values were extracted directly from Prism outputs. A total of n = 50 flies per genotype and condition were analyzed for each lifespan experiment.

Statistical analyses were conducted in GraphPad Prism, and all quantitative data are reported as mean ± SEM. Pairwise comparisons were evaluated using unpaired Student’s t-tests, whereas datasets involving multiple genotypes or conditions were analyzed using one-way ANOVA followed by either Tukey’s post-hoc testing or Dunnett’s post-hoc testing to correct for multiple comparisons. The specific statistical outcomes for each experiment are provided in the corresponding figure legends. Unless otherwise noted, significance indicators above the bars reflect comparisons to the designated control group. Values labeled ns (not significant) indicate a lack of statistical significance (p ≥ 0.05).

## Data Availability

All data supporting the findings of this study are available within the paper and its Supplementary Information. Raw images of Western blots are shown in the Supplementary Information. Additional queries pertaining to the data generated in this study can be directed to the corresponding author.

## Code Availability

Not applicable.

## Acknowledgements

We thank the Banerjee lab for critical comments on the manuscript and Priscilla McNeish for her technical assistance. We thank the NIH NeuroBioBank repository and the Harvard Brain Tissue Resource Center, for postmortem human brain tissues and Dr. Judit Oláh and Dr. Judit Ovádi for the generous gift of anti-TPPP antibodies, as well as our funding sources.

## Funding

This work was supported by funds from the National Institutes of Health/National Institute of Neurological Disorders and Stroke grants R01NS134867 and the Long School of Medicine, UT Health San Antonio. Raquel Velazquez and Michelle E. Urbina-Berlanga were part of a Summer Research Education Experience Program supported by funds from the National Institutes of Health/National Institute of Neurological Disorders and Stroke grant 5R25NS115552. The funder had no role in the design of the study, data collection, analysis, interpretation, or manuscript preparation.

## Author Contributions

**Haven Tillmon**: investigation, formal analysis, writing – original draft, writing – review and editing. **Savannah Boyen**: investigation, formal analysis, writing - review and editing. **Raquel Velazquez**: investigation, formal analysis. **Michelle E. Urbina-Berlanga**: investigation, formal analysis. **Marco Sciortino**: investigation. **Swati Banerjee**: conceptualization, project administration, resources, supervision, funding acquisition, writing – review and editing.

## Competing interests

The authors declare no competing financial or non-financial interests.

## Notes

### Competing Interest Statement

The authors have declared no competing interest.

### Summary of Updates

Major conceptual revisions include tempering claims regarding the mechanistic relationship between Ringer/TPPP dysfunction and human Parkinsons disease (PD). We revised the title and multiple sections to emphasize that our human data demonstrate a parallel rather than a causal link. We removed the previous Figure 7 and added new datasets (Figure 4C to J) showing that Ringer loss selectively impairs Complex I (CI) enzymatic activity without affecting Complexes II to V, addressing concerns about broader mitochondrial dysfunction. We also added new panels to Figure 5 (G to J) demonstrating that resveratrol does not rescue neurodegeneration, clarifying that its effects on ROS, ATP, and CI activity do not reflect restoration of underlying CI defects. A new Figure 6 presents human TPPP immunostaining in BA40, strengthening the correlative human dataset. We revised the Introduction to more clearly distinguish previously known aspects of TPPP/Ringer mitochondrial localization from the novel submitochondrial fractionation data presented here. In Figure 2, we incorporated previously supplemental CI subunit data (NDUFB8, NDUFB10, NDUFA9) to allow direct comparison of altered and unchanged proteins. We repeated and replaced the OptiPrep fractionation data in Figure 3 to present representative distributions of NDUFS2 and NDUFS3, now showing overlapping profiles consistent with their shared matrix-arm localization. We clarified limitations of density gradient fractionation and appropriately qualified mechanistic interpretations. Methodological improvements include revising CI activity quantification to subtract rotenone-inhibited values, thereby reporting true CI specific activity. We expanded ETC enzymatic analyses, corrected figure organization, and improved clarity in legends and text. We also addressed concerns regarding resveratrols broad biological effects and now interpret these findings as modulation of CI-linked oxidative stress rather than rescue of CI assembly or stability.

