## Supplementary Data for "Ringer Loss in *Drosophila* Uncovers Mitochondrial Complex I Deficits Relevant to Human Parkinson’s Disease"

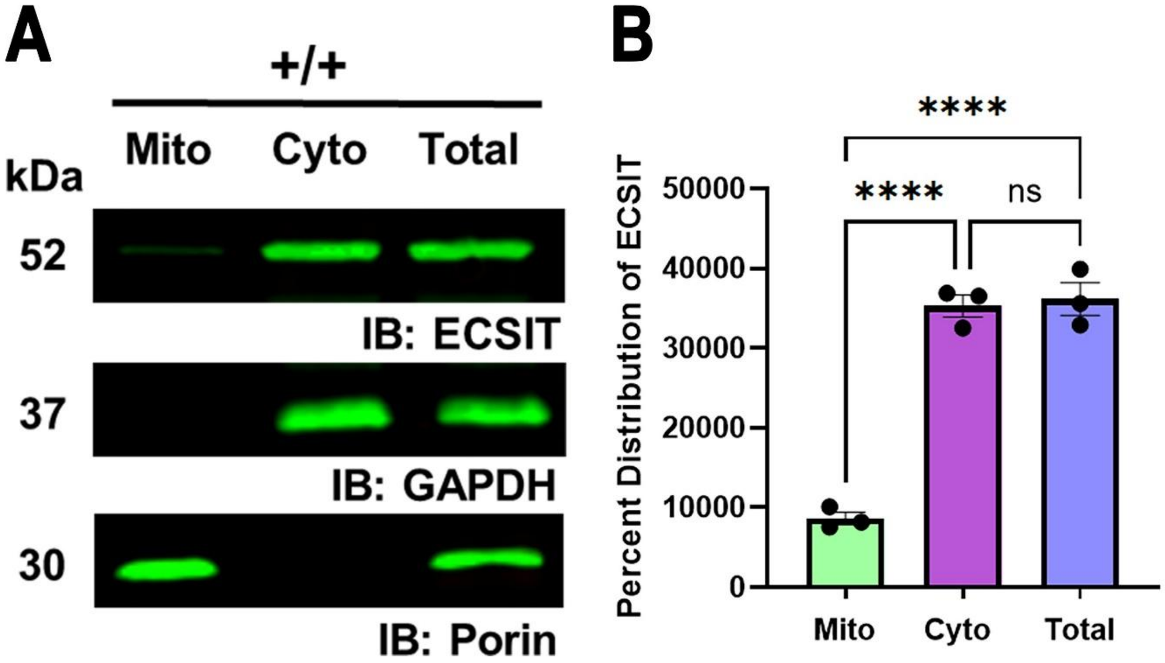


**Supplemental Figure 1: ECSIT localizes predominantly to the cytosol**

**A** Immunoblots of subcellular fractions from wild-type (+/+) fly heads probed with anti-ECSIT (52kDa), antiGAPDH (37kDa), and anti-Porin (30kDa) across mitochondrial (mito), cytosolic (cyto), and total lysate fractions. **B** Quantification of mitochondrial (green) and cytosolic (purple) band intensity as compared to the total lysate (blue) band intensity and from wild-type (+/+) fly head lysates and to each other. Data are presented as mean ±SEM. Statistical significance was determined using a one-way ANOVA followed by Tukey's multiple comparisons test. $F(6)=109.5$, ****$p<0.0001$, $n=3$.

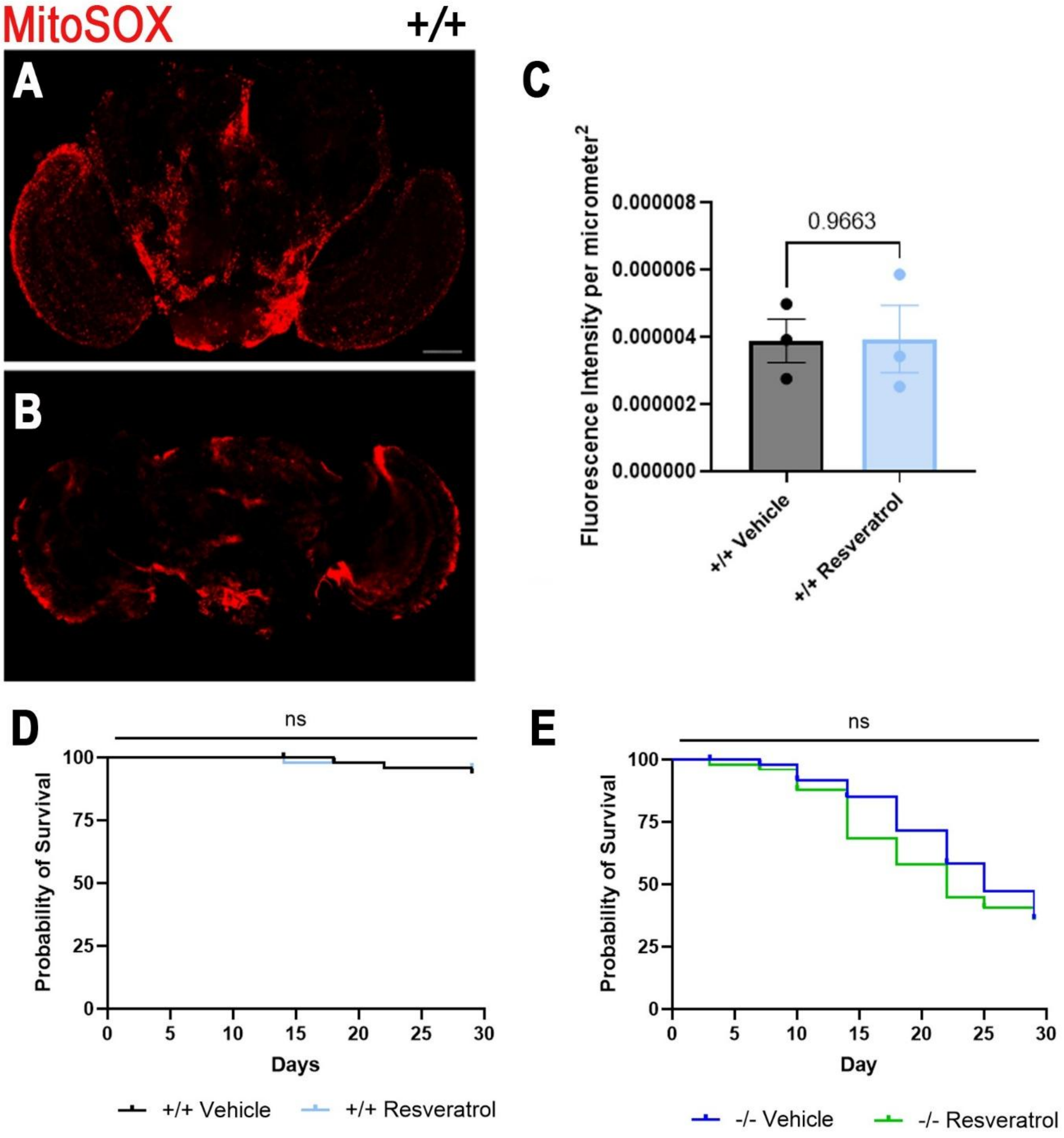


**Supplemental Figure 2: Resveratrol does not decrease ROS levels in wild-type untreated controls or alter lifespan of wild-type or ringer mutant flies** MitoSOX red fluorescence in unfixed, whole-mount adult fly brains from **A** vehicle-treated and **B** resveratrol-treated wild-type flies. **C** Quantification of MitoSOX for wild-type flies treated with vehicle (n=3) and resveratrol (n=3). Data are presented as mean ±SEM. Statistical significance was determined using a two-tailed unpaired t-test. t(4)=0.04491, ns p=0.9663. Scale bar is 100μm. Lifespan analysis of vehicle-treated and resveratrol-treated **D** wild-type (+/+) flies and **E** ringer mutant (-/-) flies. Data are presented as mean ±SEM. Statistical significance was determined using a two-tailed unpaired t-test. +/+ p=0.6942, and -/- p=0.2231. n = 50 flies per genotype.

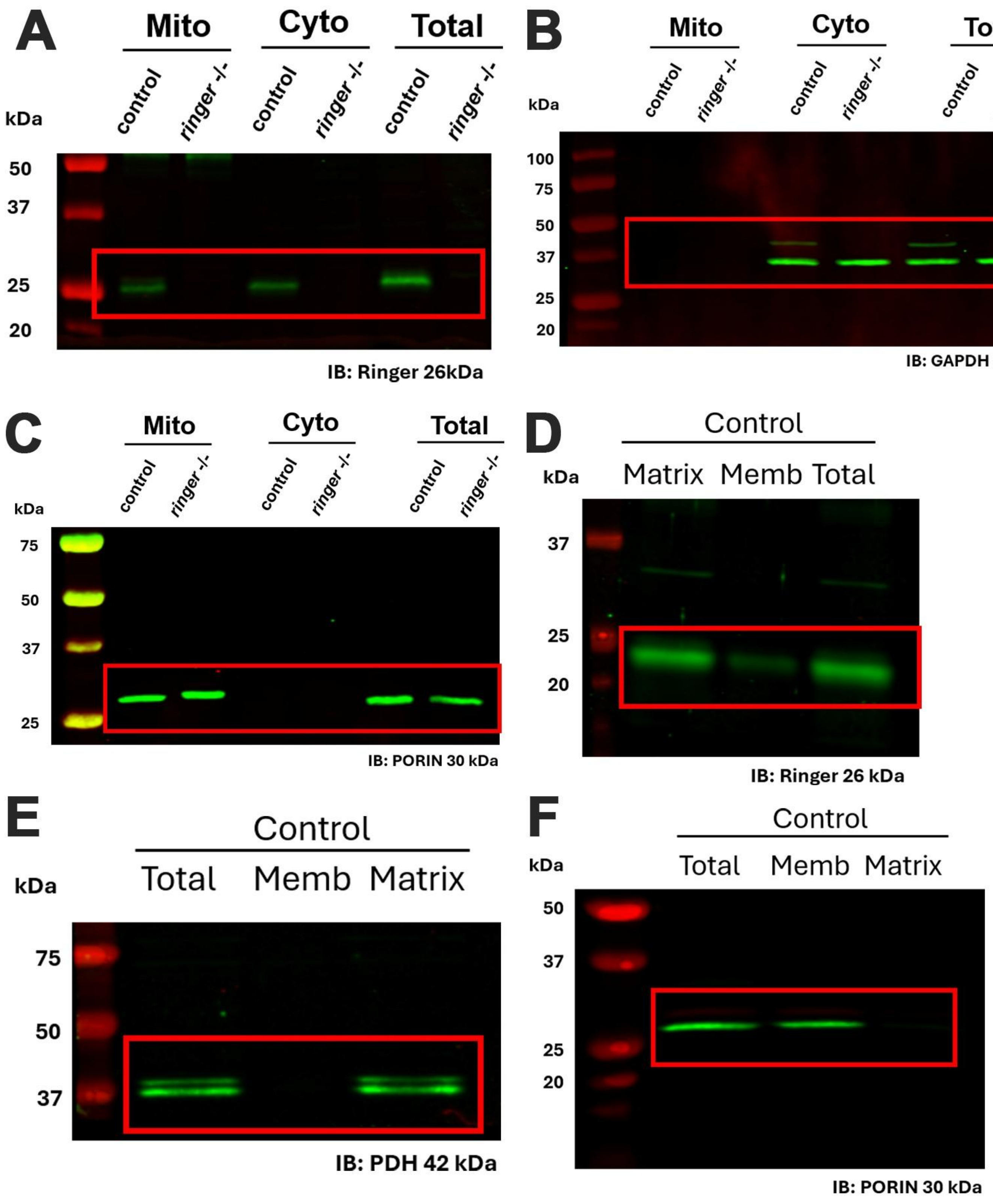
A
Mito
Cyto
Total
control
ringer -/-
control
ringer -/-
control
ringer -/-
kDa
50
37
25
20
IB: Ringer 26kDa
B
Mito
Cyto
Total
control
ringer -/-
control
ringer -/-
control
ringer -/-
kDa
100
75
50
37
25
20
IB: GAPDH 37 kDa
C
Mito
Cyto
Total
control
ringer -/-
control
ringer -/-
control
ringer -/-
kDa
75
50
37
25
IB: PORIN 30 kDa
D
Control
Matrix Memb Total
kDa
37
25
20
IB: Ringer 26 kDa
E
Control
Total Memb Matrix
kDa
75
50
37
IB: PDH 42 kDa
F
Control
Total Memb Matrix
kDa
50
37
25
20
IB: PORIN 30 kDa

**Supplemental Figure 3: Uncropped and unedited blot images corresponding to Fig. 1.** (**A-C**) Blots related to Fig. 1A. (**D-F**) Blots related to Fig. 1D.

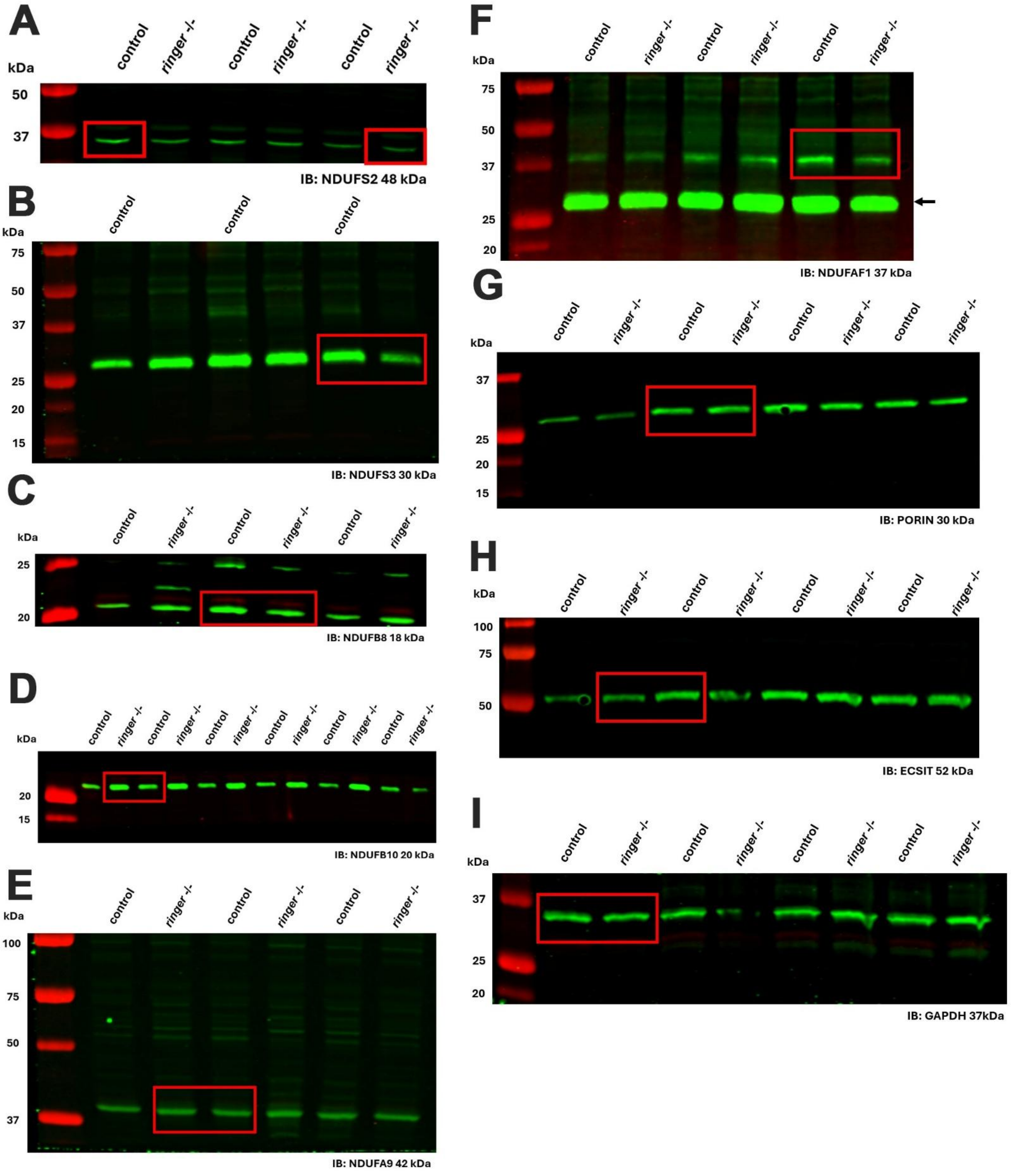
A
control
ringer -/-
control
ringer -/-
control
ringer -/-
kDa
50
37
IB: NDUFS2 48 kDa
B
control
control
control
kDa
75
50
37
25
20
15
IB: NDUFS3 30 kDa
C
control
ringer -/-
control
ringer -/-
control
ringer -/-
kDa
25
20
IB: NDUFB8 18 kDa
D
control
ringer -/-
control
ringer -/-
control
ringer -/-
control
ringer -/-
control
ringer -/-
control
ringer -/-
kDa
20
15
IB: NDUFB10 20 kDa
E
control
ringer -/-
control
ringer -/-
control
ringer -/-
kDa
100
75
50
37
IB: NDUFA9 42 kDa
F
control
ringer -/-
control
ringer -/-
control
ringer -/-
kDa
75
50
37
25
20
IB: NDUFAF1 37 kDa
G
control
ringer -/-
control
ringer -/-
control
ringer -/-
control
ringer -/-
kDa
37
25
20
15
IB: PORIN 30 kDa
H
control
ringer -/-
control
ringer -/-
control
ringer -/-
control
ringer -/-
kDa
100
75
50
IB: ECSIT 52 kDa
I
control
ringer -/-
control
ringer -/-
control
ringer -/-
control
ringer -/-
kDa
37
25
20
IB: GAPDH 37kDa

**Supplemental Figure 4: Uncropped and unedited blot images corresponding to Fig. 2.** (**A-I**) Blots related to Fig. 1A. The arrow in F refers to an unrelated antibody that the blot was probed with.

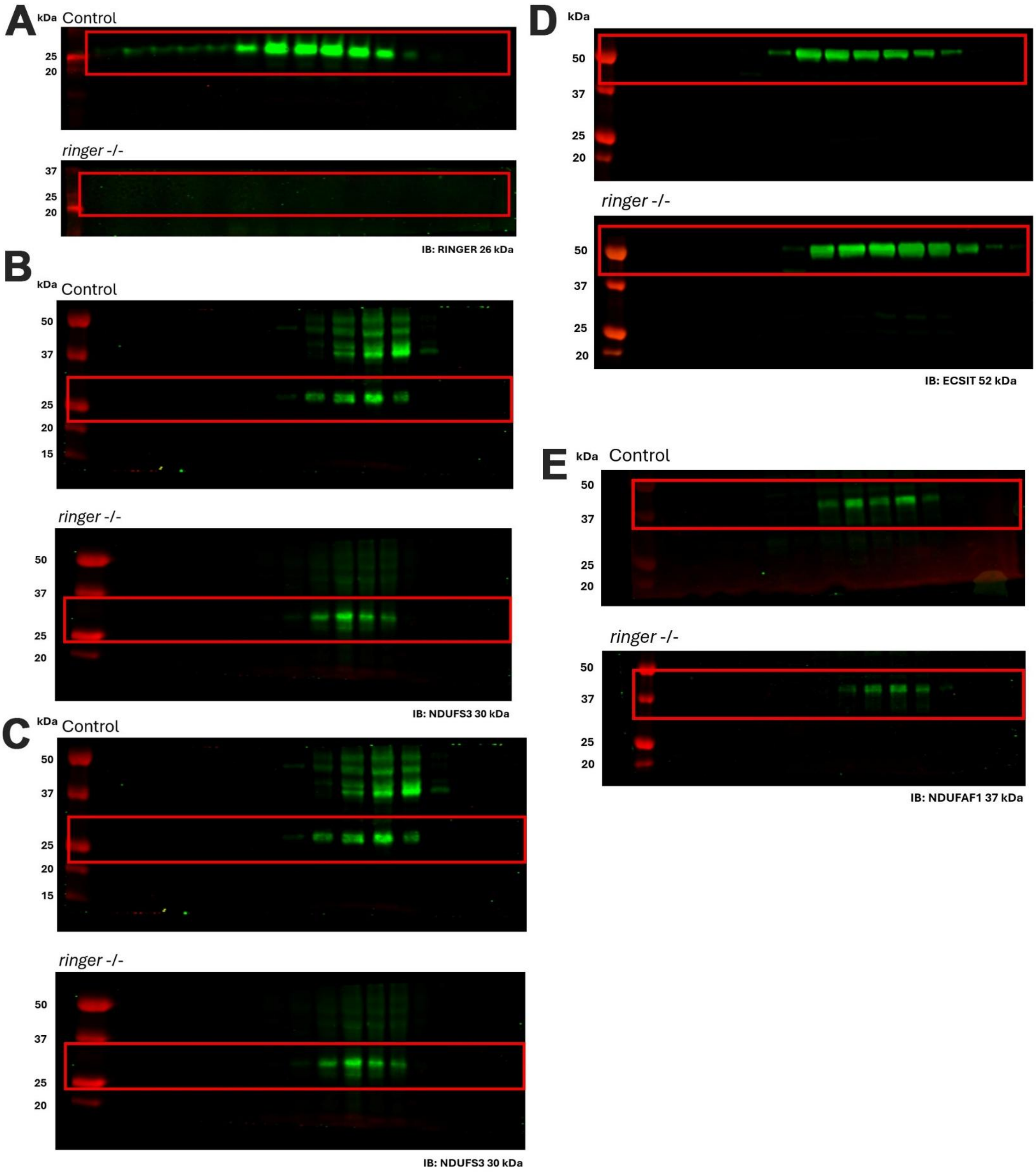
A
kDa
Control
25
20
ringer -/-
37
25
20
IB: RINGER 26 kDa
B
kDa
Control
50
37
25
20
15
ringer -/-
50
37
25
20
IB: NDUFS3 30 kDa
C
kDa
Control
50
37
25
20
15
ringer -/-
50
37
25
20
IB: NDUFS3 30 kDa
D
kDa
50
37
25
20
ringer -/-
50
37
25
20
IB: ECSIT 52 kDa
E
kDa
Control
50
37
25
20
ringer -/-
50
37
25
20
IB: NDUFAF1 37 kDa

**Supplemental Figure 5: Uncropped and unedited blot images corresponding to Fig. 3.** (A-E) Blots related to Fig. 3A-E.

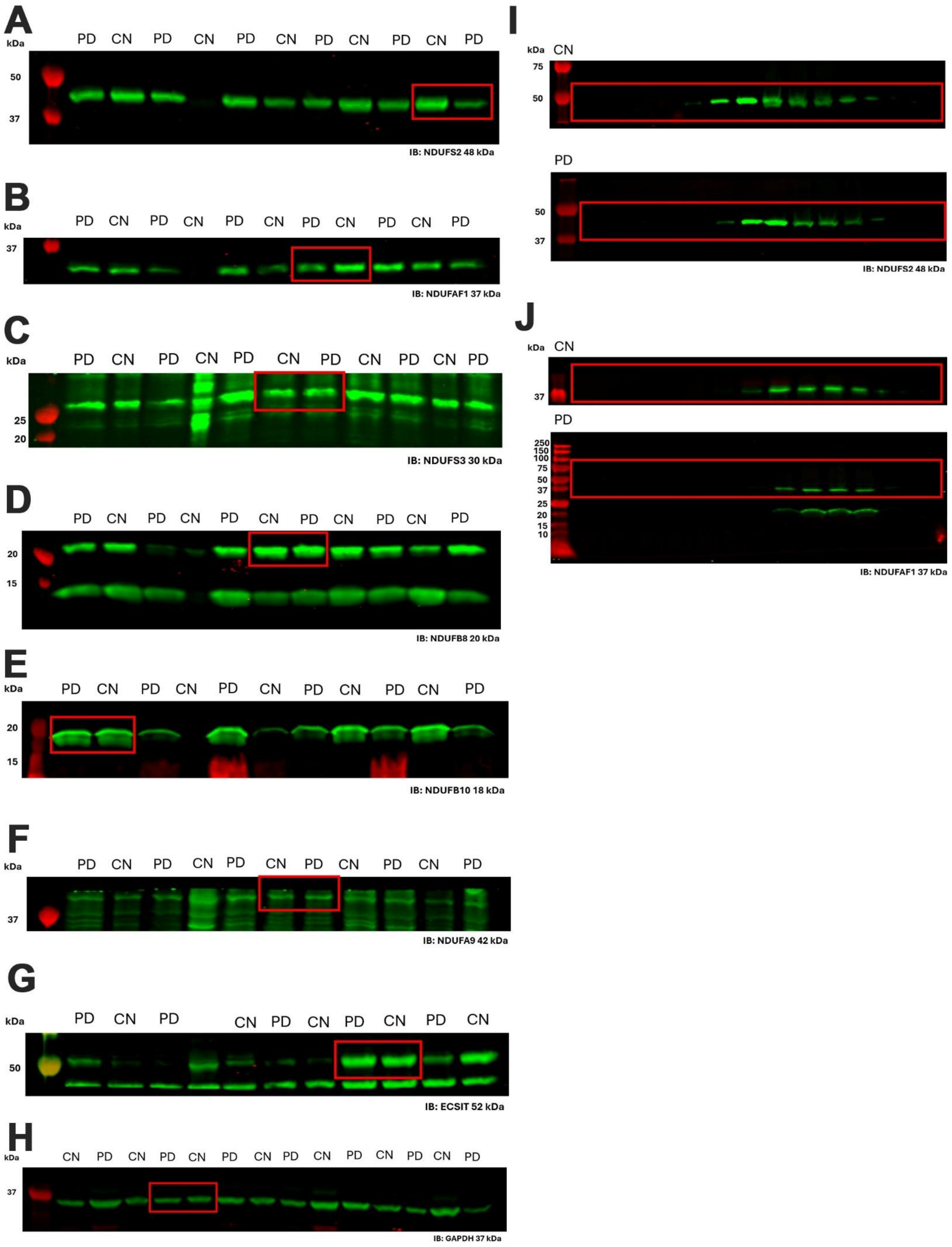
A
kDa
PD CN PD CN PD CN PD CN PD CN PD
50
37
IB: NDUFS2 48 kDa
B
kDa
PD CN PD CN PD CN PD CN PD CN PD
37
IB: NDUFAF1 37 kDa
C
kDa
PD CN PD CN PD CN PD CN PD CN PD
25
20
IB: NDUFS3 30 kDa
D
PD CN PD CN PD CN PD CN PD CN PD
20
15
IB: NDUFB8 20 kDa
E
kDa
PD CN PD CN PD CN PD CN PD CN PD
20
15
IB: NDUFB10 18 kDa
F
kDa
PD CN PD CN PD CN PD CN PD CN PD
37
IB: NDUFA9 42 kDa
G
kDa
PD CN PD CN PD CN PD CN PD CN
50
IB: ECSIT 52 kDa
H
kDa
CN PD CN PD CN PD CN PD CN PD CN PD CN PD
37
IB: GAPDH 37 kDa
I
kDa CN
75
50
PD
50
37
IB: NDUFS2 48 kDa
J
kDa CN
37
PD
250
150
100
75
50
37
25
20
15
10
IB: NDUFAF1 37 kDa

**Supplemental Figure 6: Uncropped and unedited blot images corresponding to Fig. 6.** (A-G) Blots related to Fig. 6A. (H-I) Blots related to Fig. 6I-J.
